# Coiled coil control of growth factor and inhibitor-dependent EGFR trafficking and degradation

**DOI:** 10.1101/2020.12.10.419739

**Authors:** Deepto Mozumdar, Sol Hsun-Hui Chang, Kim Quach, Amy Doerner, Alanna Schepartz

## Abstract

EGFR exhibits biased signaling, whereby growth factor or mutation-dependent changes in receptor conformation and/or dynamics elicit distinct intracellular outcomes. We report that many outcomes associated with activated EGFR are controlled by a two-state coiled coil switch located within the juxtamembrane segment (JM), an essential component of the cytosolic dimer interface. The position of this switch defines the path of endocytic trafficking and whether or not EGFR is degraded within lysosomes. JM coiled coil identity also predicts kinase-independent effects of oncogenic EGFR mutations and clinically relevant tyrosine kinase inhibitors (TKIs) that promote efficient, lysosome-based EGFR degradation. These findings provide a model for biased EGFR signaling, insights into kinase-independent activities of EGFR and clinically relevant TKIs, and identify new strategies for modulating protein lifetime.

## Introduction

Receptor tyrosine kinases in the epidermal growth factor receptor family play essential roles in cell proliferation, differentiation, and migration. Aberrant activation of EGFR family members, whether by overexpression or mutation, is associated with many human cancers. Although much work on EGFR has focused on its activation by epidermal growth factor (EGF), EGFR is activated by at least seven different growth factors including transforming growth factor α (TGF-α), epigen, epiregulin, betacellulin, heparin-binding EGF, and amphiregulin, and transmits these activation events into distinct intracellular phenotypes (Ebner and Derynck, 1991; French et al., 1995; Roepstorff et al., 2009; Ronan et al., 2016; Freed et al., 2017). Growth factor-dependent biased signaling through EGFR has been previously attributed to differences in the physical properties of the growth factors themselves (Ebner and Derynck, 1991; French et al., 1995; Freed et al., 2017) or their receptor complexes (Ronan et al., 2016; Freed et al., 2017; Macdonald-Obermann and Pike, 2014). EGFR is also activated constitutively by mutations in the kinase domain, the extracellular domain, or elsewhere (Pines et al., 2010; E. Kovacs et al., 2015); these mutations are causally linked to many human cancers (Pines et al., 2010).

Previous work has shown that the binding of most growth factors to the EGFR extracellular domain (ECD) induces the formation of one of two rotationally isomeric coiled coils within the juxtamembrane segment (JM) (Doerner et al., 2015), an essential element of the cytoplasmic dimer interface (Brewer et al., 2009; Jura et al., 2009). Binding of epidermal growth factor (EGF) or heparin-binding EGF induce a JM coiled coil defined by a leucine-rich inter-helix interface (EGF-type), whereas amphiregulin, epigen, epiregulin, or transforming growth factor α (TGF-α) induce an isomeric structure whose interface is charged and polar (TGF-α-type). JM coiled coil preference is also influenced allosterically, by point mutations within one of two transmembrane (TM) helix G-x-x-x-G motifs (Sinclair et al., 2018), oncogenic kinase domain mutations, and, in the case of drug-resistant L834R/T766M EGFR, kinase domain pharmacologic status (Lowder et al., 2015).

While it is clear that the JM segment is coupled allosterically to the ECD, TM, and kinase domains, this very coupling complicates strategies to define how and whether distinct JM coiled coil structures influence outcomes associated with biased signaling. To better understand the relationship between JM coiled structure and EGFR biology, we designed a set of kinase-active EGFR mutants that effectively shift the equilibrium between JM coiled coil isomers. These EGFR mutants assemble into dimers that favor one coiled coil or the other in a manner that is independent of the activating growth factor. Using these and other “decoupling” mutants, we demonstrate that coiled coil identity alone is necessary and sufficient to define the pathway of EGFR trafficking into degradative (Rab7+) or recycling (Rab11+) endosomes and ultimately whether EGFR is degraded within lysosomes. JM coiled coil identity also predicts the trafficking and lifetime of oncogenic EGFR mutants and reveals kinase-independent effects of FDA-approved small molecule tyrosine kinase inhibitors (TKIs). These discoveries increase our understanding of the molecular mechanism used for biased signaling in ErbB-family receptors and suggest new strategies for purposefully controlling protein traffic and lifetime along the endocytic pathway.

## Results

### Design of EGFR decoupling mutants

First we sought to identify a set of functional, full-length EGFR variants that isolated the cellular role of the JM by decoupling the well-established relationship between JM coiled coil status and growth factor identity (Doerner et al., 2015; Sinclair et al., 2018; Scheck et al., 2012). It is well known that salt bridges at the *e* and *g* positions of model coiled coils influence stability and orientation (O’Shea et al., 1993), while changes at the *a* and *d* positions influence oligomeric state (Harbury et al., 1993). These studies suggested that the relative stability of the EGF- and TGF-α-type coiled coils formed within an intact EGF receptor dimer could be controlled by the presence or absence of coiled coil-specific salt-bridging residues at positions *e* and *g* (Figure. 1a). We hypothesized that removing one or more interactions that stabilized one JM coiled coil would raise its free energy relative to the other and shift the equilibrium between the two structures (Figure 1–figure supplement 1a). The result would be a set of EGFR variants that favored one and only one coiled coil, regardless of the identity of the activating growth factor. The EGF- and TGF-α-type coiled coils are equally populated when no growth factor-dependent signal emerges from the extracellular domain (Mozumdar et al., 2020).

**Figure. 1.**
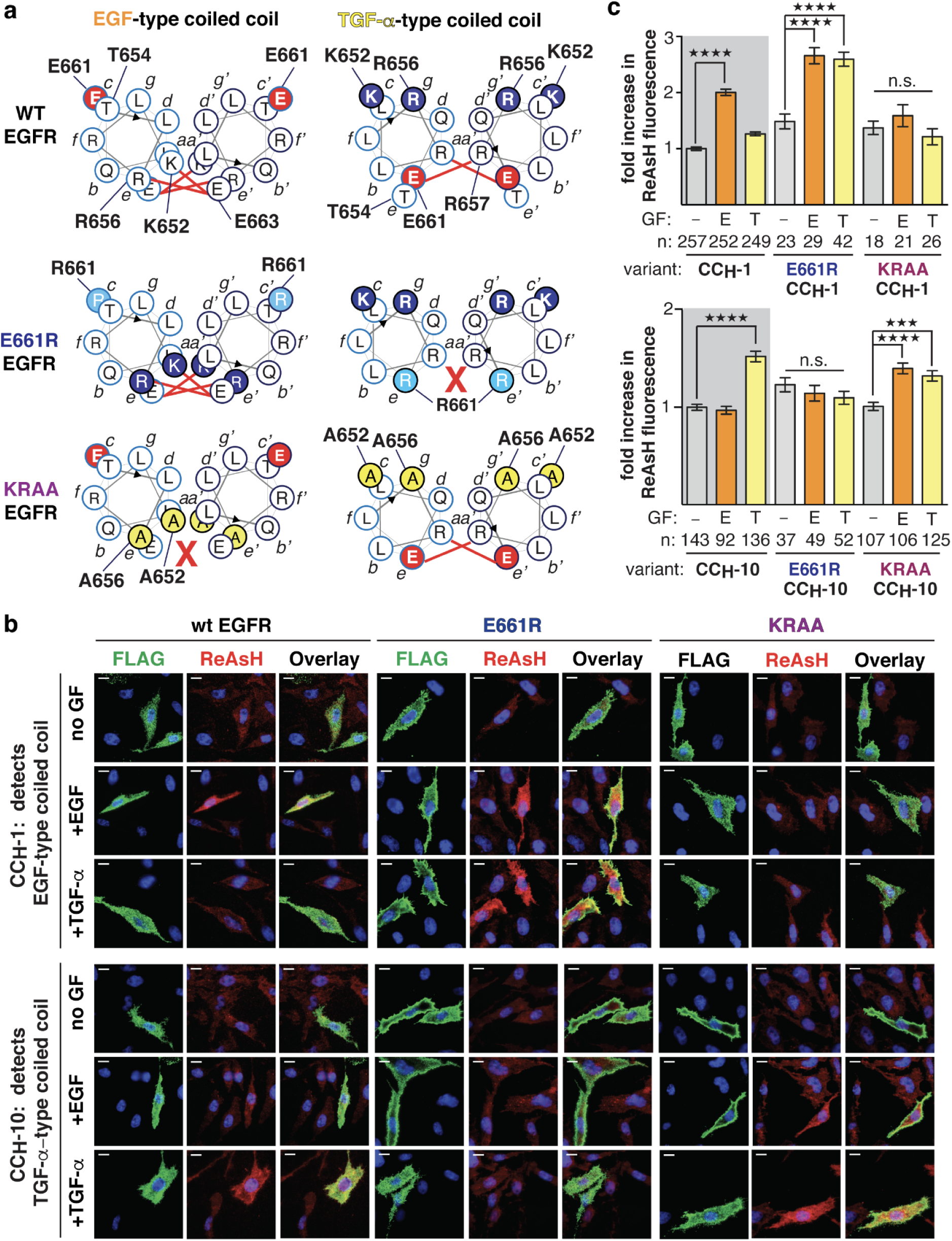
Design of E661R and KRAA EGFR decoupling mutants. **(a)** Helical wheel diagrams illustrating relative positions of residues within the N-terminal region of the EGFR JM segment of WT, E661R, and KRAA EGFR when assembled into an EGF-type (left) or TGF-α-type (right) coiled-coil. Bold red lines identify potential salt bridge interactions referred to in the text. **(b)** Representative TIRF-M images of CHO-K1 cells illustrating ReAsH labeling (red fluorescence) and expression (green fluorescence) of FLAG-tagged CC_H_-1 and CC_H_-10 variants of WT, E661R, or KRAA EGFR with or without EGF or TGF-α stimulation (16.7 nM). Scale bars = 10 µm. **(c),** Bar plots illustrating the fold increase in expression-corrected ReAsH fluorescence over background. n = # of cells. Error bars = s.e.m. ****p<0.0001, ***p<0.001, **p<0.01, *p<0.1 from one-way ANOVA with Bonferroni post-analysis accounting for multiple comparisons. n.s., not significant. Figure 1–figure supplement 1 shows representative TIRF-M images of CHO-K1 cells illustrating ReAsH labeling (red fluorescence) and expression (green fluorescence) of FLAG-tagged CC_H_-1 and CC_H_-10 variants of T654D EGFR with or without EGF or TGF-α stimulation (16.7 nM). and data for expression and activity of FLAG-tagged CC_H_-1 and CC_H_-10 variants of WT, E661R, KRAA and T654D EGFR.

To test this hypothesis, we generated a pair of EGFR variants (E661R and KRAA) containing one or two amino acid substitutions that selectively disrupt salt bridges unique to either the TGF-α- or EGF-type JM coiled coils (Figure 1a and Figure 1–figure supplement 1b). The single charge-reversing mutation in E661R EGFR is located at distal *c* and *c’* positions of the heptad repeat when the JM folds into an EGF-type coiled coil, but at proximal *e* and *e’* positions when it folds into a TGF-α-type structure. As a result, the JM in E661R EGFR should favor the EGF-type structure because it lacks a TGF-α-type-specific salt-bridge. By contrast, the two charge-eliminating substitutions in K652A/R656A EGFR (KRAA) are located at distal *c,g* and *c’,g’* positions of the coiled coil repeat when the JM is assembled into the TGF-α-type structure, but at proximal *e* and *a* positions when assembled into the EGF-type structure. The JM in KRAA EGFR should therefore favor the TGF-α-type structure because it lacks two EGF-type-specific salt bridges (Figure 1a). T654D EGFR was prepared as a control: this mutation occupies the *c* or *e* position of a coiled coil repeat when the JM assembles into the EGF- and TGF-α-type structures, respectively (Figure 1a). All variants could be expressed in CHO-K1 cells (which express little or no endogenous EGFR (Krug et al., 2003)), trafficked to the cell surface, and were phosphorylated at multiple tyrosine residues within the C-terminal tail when treated with saturating (16.7 nM) EGF or TGF-α (Figure 1– figure supplement 1c).

### Validating EGFR decoupling mutants

With functional mutant EGF receptors in hand, we made use of a chemical biology tool, bipartite tetracysteine display (Scheck et al., 2012), to test the hypothesis that activated E661R and KRAA EGFR dimers would favor an EGF-type or TGF-α-type coiled coil, respectively, regardless of the identity of the activating growth factor. Bipartite tetracysteine display exploits the pro-fluorescent bis-arsenical dye ReAsH (Adams et al., 2002), which lights up only when bound to four Cys side chains (two from each EGFR monomer) in a defined array (Walker et al., 2016). Previous work identified a CysCys-containing EGFR variant whose dimer induces ReAsH fluorescence only when assembled into an EGF-type type JM coiled coil (EGFR CC_H_-1); another CysCys-containing EGFR variant (EGFR CC_H_-10) forms dimers that induce ReAsH fluorescence only when the TGF-α-type coiled coil is formed (Doerner et al., 2015). Variants of CC_H_-1 and CC_H_-10 EGFR containing E661R or KRAA mutations were phosphorylated at multiple tyrosine residues within the C-terminal tail when treated with EGF or TGF-α; EGFR T654D was somewhat less active (Figure 1–figure supplement 1d).

The three sets of EGFR CC_H_-1 and CC_H_-10 variants were expressed in CHO-K1 cells, stimulated with EGF or TGF-α, incubated with ReAsH, and the level of ReAsH fluorescence relative to EGFR-expression determined using TIRF microscopy (TIRF-M) (Figure 1b and c). TIRF-M excites fluorophores in an extremely thin axial region, typically within ∼100 nm of the cell surface, thus the measured fold-increases in ReAsH fluorescence provide a read-out of JM coiled coil conformation within EGFR molecules at or near the plasma membrane. As expected (Doerner et al., 2015), a significant fold-increase in ReAsH fluorescence was observed when cells expressing WT EGFR CC_H_-1 EGFR are treated with EGF (2.00 ± 0.06) but not TGF-α (1.26 ± 0.03), or when cells expressing WT CC_H_-10 EGFR were treated with TGF-α (1.52 ± 0.05) and not EGF (0.97 ± 0.04). By contrast, cells expressing E661R CC_H_-1 EGFR showed a significant fold-increase in ReAsH fluorescence regardless of whether the cells were treated with EGF (2.66 ± 0.14) or TGF-α (2.60 ± 0.12), suggesting that the EGF-type structure formed in both cases. Little or no fold-increase in ReAsH fluorescence was observed when cells expressing E661R CC_H_-10 EGFR were treated with EGF (1.14 ± 0.08) or TGF-α (1.10 ± 0.07).

In a similar way, cells expressing KRAA CC_H_-10 EGFR displayed a significant fold-increase in ReAsH fluorescence regardless of whether the cells were treated with EGF (1.40 ± 0.05) or TGF-α (1.32 ± 0.05), and little increase was observed in cells expressing KRAA CC_H_-1 EGFR (1.59 ± 0.19-fold (EGF); 1.21 ± 0.14-fold (TGF-α)) (Figure 1b and c). Analogous variants containing T654D mutations behaved like WT EGFR, as expected (Figure 1–figure supplement 1e and f) and no fold-increase in ReAsH fluorescence was observed in the absence of added growth factor. These data indicate that no matter which growth factor is bound to the EGFR extracellular domain, E661R EGFR assembles into an active dimer containing an EGF-type coiled coil, whereas KRAA EGFR assembles into an active dimer containing a TGF-α-type coiled coil. These results confirm that the mutations embodied by E661R and KRAA EGFR effectively decouple growth factor identity from coiled coil status.

### Trafficking of E661R and KRAA EGFR

A quintessential growth factor-dependent EGFR activity is the path of receptor trafficking following endocytosis (Bakker et al., 2017). Unactivated EGF receptors at the cell surface are internalized into Rab4-associated early endosomes and recycled back to the cell surface where they accumulate. Activated receptors are trafficked first to EEA1-positive (EEA1+) early endosomes (Mu et al., 1995). The pathway then splits, and receptors are sorted into vesicles defined by the presence of either Rab11 (Rab11+) or Rab7 (Rab7+). The former deliver EGFR back to the cell surface (recycling pathway) (Ullrich et al., 1996) whereas the latter deliver EGFR to late endosomes and lysosomes where the receptors are ultimately degraded (degradative pathway) (Ceresa and Bahr, 2006). Trafficking from EEA1+ early endosomes into recycling or degradative endosomes is growth factor-dependent: when stimulated with EGF, EGFR traffics into Rab7+ endosomes and is ultimately degraded (Ceresa and Bahr, 2006); when stimulated with TGF-α, EGFR traffics into Rab11+ endosomes and returns to the cell surface (Francavilla et al., 2016).

The dependence of the path of EGFR trafficking on growth factor identity has been previously attributed to differences in the pH-dependent ligand occupancy of endocytosed receptors (Ebner and Derynck, 1991; French et al., 1995). In particular, TGF-α dissociates from EGFR at higher pH than does EGF; ligand dissociation at earlier points along the endocytic pathway is believed to lead ultimately to receptor sorting. Although the pH-dependent model is simple, it is inconsistent with reports that TGF-α-bound EGFR continues to signal within endosomes (Francavilla et al., 2016). We thus asked whether JM coiled coil status could also control the pathway of EGFR trafficking. If so, then EGFR mutations that favor the EGF-type (E661R) coiled coil in activated dimers would bias trafficking into Rab7+ endosomes (degradative pathway), whereas those that favor the TGF-α-type structure would bias trafficking into Rab11+ endosomes (recycling pathway).

We used confocal microscopy to trace the time-, mutation, and growth factor-dependent pattern of EGFR trafficking and organelle co-localization following growth factor stimulation. CHO-K1 cells expressing FLAG-tagged WT, E661R, KRAA or T654D EGFR were incubated first on ice with EGF or TGF-α to allow growth factor binding and then at 37 °C in growth factor-free media to initiate endocytosis and receptor trafficking. After 8 or 40 min, the cells were immuno-stained to visualize EGFR and assess the extent of colocalization with EEA1, Rab7, or Rab11. WT EGFR, E661R, KRAA, and T654D EGFR all co-localize with the early endosome marker EEA1 and not with Rab7 or Rab11 after 8 min (Figure 2–figure supplement 1), regardless of whether the cells were stimulated with EGF or TGF-α. After 40 min, the colocalization of all activated receptors with EEA1 decreases to non-significant levels (Roepstorff et al., 2009) (Figure 2–figure supplement 2a and b). These results confirm that the growth factor-dependent trafficking of all EGFR variants studied here proceeds initially through EEA1+ early endosomes regardless of JM coiled coil status or growth factor identity.

**Figure 2.**
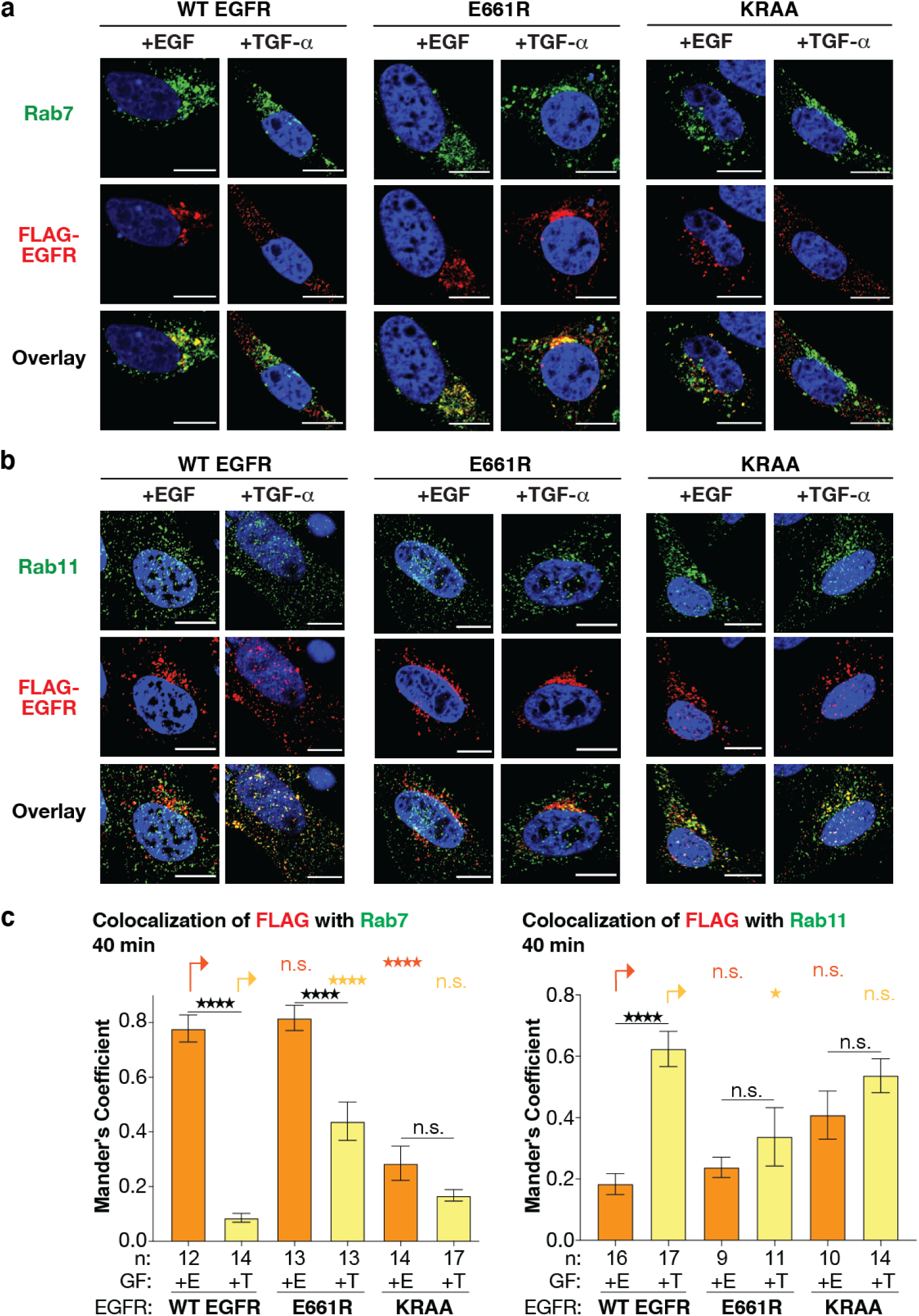
The path of EGFR trafficking in CHO-K1 cells is controlled by JM coiled coil identity. Confocal microscopy of CHO-K1 cells expressing FLAG-tagged WT, E661R, or KRAA EGFR (false colored red), immuno-labeled with **(a)** Rab7 (false colored green) as a marker for degradative endosomes or **(b)** Rab11 (false colored green) as a marker for recycling endosomes, 40 minutes after stimulation with EGF (E) or TGF-α (T). Scale bars = 10 µm. **(c)** Bar plots illustrating the Manders colocalization coefficient (MCC) of FLAG-tagged WT, E661R and KRAA EGFR with either Rab7 or Rab11 40 minutes after stimulation with EGF/TGF-α. n = # of cells. Error bars = s.e.m. ****p<0.0001, ***p<0.001, **p<0.01, *p<0.1, n.s. not significant, from one-way ANOVA with Tukey’s multiple comparison test. See also Figure 2– figure supplement 1, Figure 2– figure supplement 2 for trafficking controls, Figure 2 – figure supplement 3 for time dependent decay of phosphorylation of JM mutants.

We used the same workflow to evaluate the post-EEA1+ trafficking pattern of FLAG-tagged WT, E661R, KRAA and T654D EGFR. WT EGFR and T654D EGFR both colocalized preferentially with Rab7 when activated with EGF (MCC = 0.78 ± 0.05 (WT) and 0.71 ± 0.05 (T654D)) and with Rab11 when activated with TGF-α (MCC = 0.62 ± 0.06 (WT) and 0.60 ± 0.06 (T654D)) (Figure 2 and Figure 2–figure supplement 2c and d). In contrast, while E661R still colocalized significantly with Rab7 when activated with EGF, it also colocalized significantly with Rab7 when activated with TGF-α. The extent of EGFR colocalization with Rab7 upon TGF-α activation was low (MCC = 0.09 ± 0.02) for WT EGFR but moderate (MCC = 0.44 ± 0.07) for the E661R variant (Figure 2). Similarly, the extent of colocalization with Rab11 upon TGF-α activation was high (MCC = 0.62 ± 0.06) for WT EGFR but only moderate (MCC = 0.34 ± 0.10) for the E661R variant. These results indicate that the E661R mutation biases EGFR trafficking into Rab7+ endosomes regardless of whether EGF or TGF-α is used to activate the receptor.

The inverse set of results were obtained when the pathway of KRAA EGFR trafficking was examined: KRAA EGFR colocalized significantly with Rab11 (and not Rab7) whether activated with EGF or TGF-α. The extent of EGFR colocalization with Rab11 upon EGF activation was low (MCC = 0.18 ± 0.03) for WT EGFR but moderate (MCC = 0.41 ± 0.08) for the KRAA variant (Figure 2). Similarly, the extent of colocalization with Rab7 upon EGF activation was high (MCC = 0.78 ± 0.05) for WT EGFR but only moderate (MCC = 0.29 ± 0.06) for the KRAA variant. These results indicate that the KRAA mutation biases EGFR trafficking into Rab11+ endosomes regardless of whether EGF or TGF-α is used to activate the receptor. Taken together, these results indicate that the pathway of EGFR trafficking is influenced by the structure formed within the juxtamembrane segment that links the TM region to the kinase domain. We note, however, that the biasing of EGFR trafficking is not a perfect “on/off” switch: while the E661R substitution increases the fraction of EGFR that traffics into Rab7+ endosomes in the presence of TGF-α, a small fraction of TGF-α-activated E661R EGFR traffics into Rab11+ endosomes. Indeed, the extent of phosphorylation at C-terminal tail residues Y1045, Y1068, and Y1173 did not obviously correlate with JM mutational state (Figure 2–Figure Supplement 3a-c).

**Figure 3:**
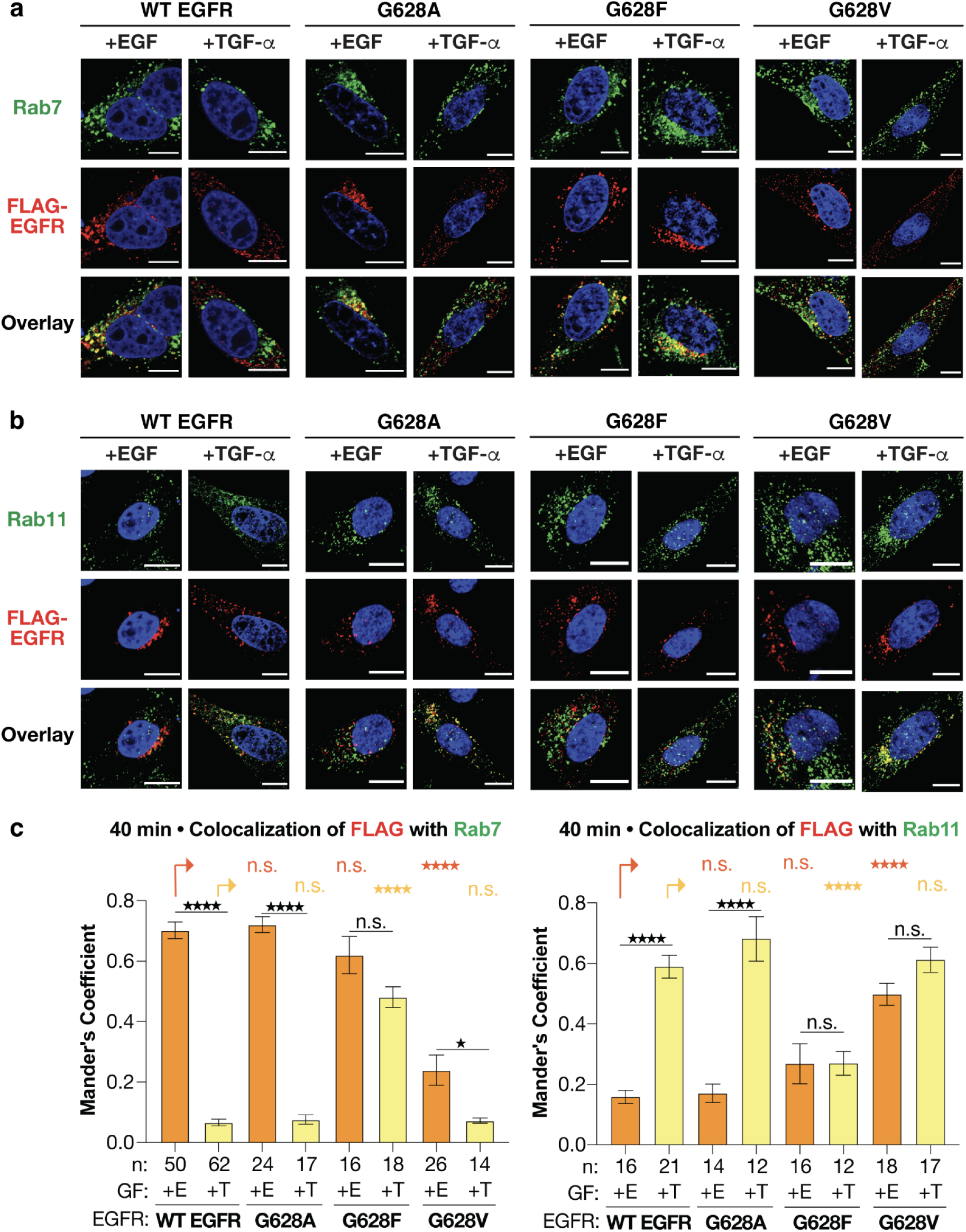
Point mutations within the EGFR transmembrane helix allosterically influence the pathway of receptor trafficking. Confocal microscopy of CHO-K1 cells expressing FLAG-tagged G628A, G628F and G628V EGFR (false colored red) and immuno-labeled with **(a)** Rab7 (false colored green) as a marker for degradative endosomes or **(b)** Rab11 (false colored green) as a marker for recycling endosomes, 40 minutes after stimulation with EGF (E) or TGF-α (T). Scale bars = 10 µm. **(c)** Bar Plots illustrating colocalization of FLAG-tagged G628A, G628F and G628V EGFR with either Rab7 or Rab11-GFP 40 minutes after stimulation with EGF/TGF-α (16.7 nM). n = # of cells. Error bars = s.e.m.. ****p<0.0001, ***p<0.001, **p<0.01, *p<0.1, n.s. not significant from one-way ANOVA with Tukey’s multiple comparison test. See also Figure 3–figure supplement 1, for trafficking controls, Figure 3–figure supplement 2 for time dependent decay of phosphorylation of TM mutants.

### Mutations within the EGFR transmembrane helix that allosterically influence JM coiled coil status control the pathway of receptor trafficking

Mindful of the fact that E661R and KRAA EGFR contain mutations within the JM coiled coil itself which could influence functional intra- or intermolecular protein interactions, we turned to a new set of EGFR decoupling variants in which JM coiled coil identity is controlled allosterically by point mutations in the adjacent transmembrane segment (TM) (Sinclair et al., 2018).

Substitution of phenylalanine for a single glycine (G628) at the final position of the N-terminal G-x-x-G motif of the EGFR TM (G628F EGFR) generates a receptor dimer with an EGF-type JM coiled coil, regardless of whether the receptor is activated by EGF or TGF-α–just like E661R EGFR. Likewise, substitution of the beta-branched residue valine at position 628 (G628V EGFR) generates a receptor dimer with a TGF-α-type coiled coil, regardless of whether the receptor is activated by EGF or TGF-α–just like KRAA EGFR (Sinclair et al., 2018). Substitution of G628 with alanine (G628A EGFR) generates a receptor that behaves like WT EGFR. Since these TM variants contain a wild-type JM, their functional intra- or intermolecular protein interactions should be preserved. As a result, the trafficking patterns they follow provide an unadulterated view of the relationship between JM conformational status and EGFR trafficking. If JM coiled coil structure is both necessary and sufficient to direct the pathway of EGFR trafficking, then G628F and G628V EGFR should traffic into only Rab7+ or Rab11+ endosomes, respectively, regardless of how they are activated, whereas G628A EGFR should behave like WT EGFR and traffic in a growth factor-dependent manner. Control experiments verified that G628F, G628V, and G628A EGFR colocalize after 8 min with EEA1 (Figure 3–figure supplement 1a and b) and not with Rab7 (Figure 3–figure supplement 1c and d) or Rab11 (Figure 3–figure supplement 1e and f); colocalization with EEA1 falls to expected levels after 40 min (Figure 3–figure supplement 1g and h).

Indeed, after 40 minutes, although the extent of G628A EGFR co-localization with Rab7 and Rab11 was growth factor-dependent, the localization of G628F and G628V EGFR were not (Figure 3a and b). When stimulated with EGF, G628A EGFR co-localized preferentially with Rab7 (MCC = 0.38 ± 0.06) and not Rab11 (MCC = 0.13 ± 0.03), when stimulated with TGF-α, G628A EGFR co-localized preferentially with Rab11 (MCC = 0.33 ± 0.03) and not Rab7 (MCC = 0.13 ± 0.03). By contrast, G628F EGFR colocalizes exclusively with Rab7 whether stimulated with EGF (MCC = 0.48 ± 0.05) or TGF-α (MCC = 0.37 ± 0.03) and not with Rab11 (MCC = 0.11 ± 0.05 and 0.14 ± 0.04 for cells stimulated with EGF and TGF-α-, respectively). Conversely, G628V EGFR colocalizes exclusively with Rab11 whether stimulated with EGF (MCC = 0.65 ± 0.03) or TGF-α (MCC = 0.65 ± 0.07) and not with Rab7 (MCC = 0.12 ± 0.02 and 0.16 ± 0.03 for cells stimulated with EGF and TGF-α-, respectively). Here, when JM coiled coil status is determined by distal mutations, the switch in trafficking pattern was complete: G628F EGFR, whose JM contains an EGF-type coiled coil, trafficks along the degradative pathway whereas G628V EGFR, whose JM contains a TGF-type coiled coil, trafficks along the recycling pathway. In this case, the extent of phosphorylation at C-terminal tail residues Y1045,Y1068, and Y1173 correlates with JM mutational state (Figure 3–figure supplement 2).

### Coiled coil control of EGFR degradation

The differences in endocytic trafficking seen with EGFR TM mutants that influence JM structure result in predictable changes in EGFR lifetime (Figure 4a and b). The lifetime of WT and G628A EGFR depends on growth factor identity in the expected way (Francavilla et al., 2016; Roepstorff et al., 2009). Following EGF stimulation, WT and G628A EGFR levels decrease rapidly and the fraction of intact receptor detected after 90 minutes is low (41-45%), whereas the fraction of intact EGFR detected after 90 minutes is high when cells are stimulated with TGF-α (94-92%). By contrast, G628F EGFR is degraded rapidly regardless of whether the receptor was activated with EGF or TGF-α, with 42% and 60% of the intact receptor remaining after 90 min following EGF or TGF-α treatment, respectively (Figure 4a and b). Conversely, G628V was degraded slowly following treatment with EGF or TGF-α, with 92% and 95% of the intact receptor remaining after 90 min, respectively (Figure 4a and b). Control experiments confirmed that EGFR degradation occurred within lysosomes (Figure 4a and b). Thus, the presence of a single point mutation in the TM segment and its long-range effect on JM conformation are necessary and sufficient to dictate the pathway of intracellular trafficking and the extent of EGFR degradation in lysosomes.

**Figure 4.**
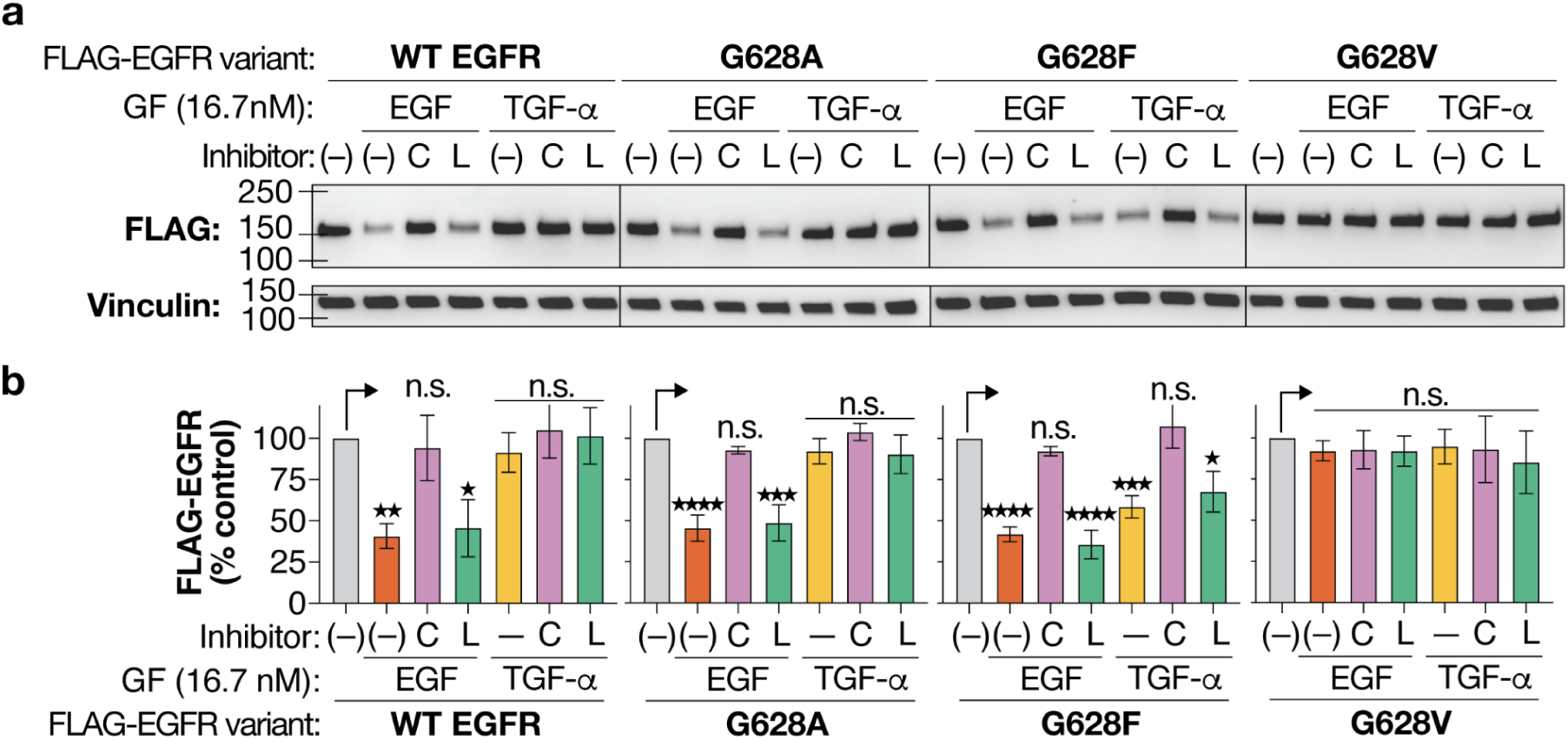
Coiled coil control of EGFR degradation. **(a)** Immunoblots illustrating the level of FLAG-tagged WT, G628A, G628F, and G628V EGFR detected in CHO-K1 cells 90 minutes after stimulation with or without EGF/TGF-α (16.7 nM) and without/with pre-incubation with the lysosomal inhibitor chloroquine, C (100 µM)(Homewood et al., 1972) or the proteasomal inhibitor lactacystin, L (10 µM)(Fenteany et al., 1995) for 1 hour at 37 °C. **(b)** Plot illustrating the normalized percent of intact FLAG-tagged WT, G628A, G628F, and G628V EGFR as shown in **4a**. Vinculin is used as loading control. Error bars = s.e.m.. ****p<0.0001, ***p<0.001, **p<0.01, *p<0.1, n.s. not significant from one-way ANOVA with Dunnett’s multiple comparison test.

### Tyrosine kinase inhibitors influence the trafficking path and lifetime of L834R/T766M EGFR

The allosteric network that coordinates information transfer between the ECD, TM, and JM regions of EGFR also includes the cytoplasmic kinase domain and its pharmacologic state (Lowder et al., 2015; Sinclair et al., 2018; Macdonald-Obermann and Pike, 2018). Kinase domain mutations associated with drug-resistant non-small cell lung cancer (L834R/T766M) (Lynch et al., 2004) generate constitutively active receptors whose JM favors a TGF-α-type structure (Lowder et al., 2015). The structure of the JM coiled coil shifts into the EGF-type structure when L834R/T766M EGFR is inhibited by selective third-generation tyrosine kinase inhibitors (TKIs) such as WZ-4002 (Zhou et al., 2009), CO-1686 (Walter et al., 2013), or the clinically approved AZD-9291 (Cross et al., 2014). Previous work has shown that L834R/T766M EGFR is constitutively endocytosed and recycled in H1975 cells (Chung et al., 2009), as predicted by the position of its JM coiled coil switch. Here we ask whether this trafficking pattern is also observed in CHO-K1 cells expressing L834R/T766M EGFR, whether it is affected by L834R/T766M EGFR-selective TKIs, and whether differences in trafficking lead to predictable changes in L834R/T766M EGFR lifetime.

Using confocal microscopy, we evaluated the pattern of endocytic trafficking and sub-cellular localization of L834R/T766M EGFR in CHO-K1 cells in the presence and absence of second- (Afatinib), third- (AZD-9291, CO-1686, WZ-4002), and fourth-generation (EAI045) TKIs (Jia et al., 2016). These small molecule EGFR inhibitors differ in mechanism of engagement (covalent *vs.* non-covalent), EGFR specificity (WT *vs.* DM; monomer *vs.* dimer-specific), and binding site (ATP *vs.* allosteric) (Figure 5). All decreased the levels of EGFR auto-phosphorylation (Figure 5–figure supplement 1a). As anticipated from data in H1975 cells (Chung et al., 2009), after 40 min uninhibited L834R/T766M EGFR colocalizes with Rab11 (MCC = 0.48 ± 0.03) and not Rab7 (MCC = 0.17 ± 0.01) (Figure 5a-d), favoring the recycling pathway expected for activated receptors that contain a TGF-α-type JM coiled coil (Lowder et al., 2015). Afatinib-inhibited L834R/T766M EGFR also colocalizes with Rab11 (MCC = 0.46 ± 0.04) and not Rab7 (MCC = 0.14 ± 0.02) after 40 min, again favoring the recycling pathway expected for activated receptors that contain a TGF-α-type JM coiled coil (Lowder et al., 2015). By contrast, when covalently inhibited by AZD-9291, CO-1686, or WZ-4002, all third-generation TKIs that selectively and covalently inhibit L834R/T766M EGFR, the extent of L834R/T766M EGFR colocalized with Rab7 increases and colocalization with Rab11 decreases (Figure 5b and d). Notably, these changes are not observed in cells expressing L834R/T766M EGFR and treated with the fourth-generation inhibitor EAI045, which binds noncovalently in a pocket adjacent to that of AZD-9291 (Jia et al., 2016). EAI045 differs from AZ-9291, CO-1686, and WZ-4002 in both engagement mode and binding site within the EGFR kinase domain, suggesting that the observed differences in receptor trafficking likely result from a common allosteric change in EGFR structure induced by these three small molecules that guides the receptor along the degradative arm of the endocytic pathway.

**Figure 5.**
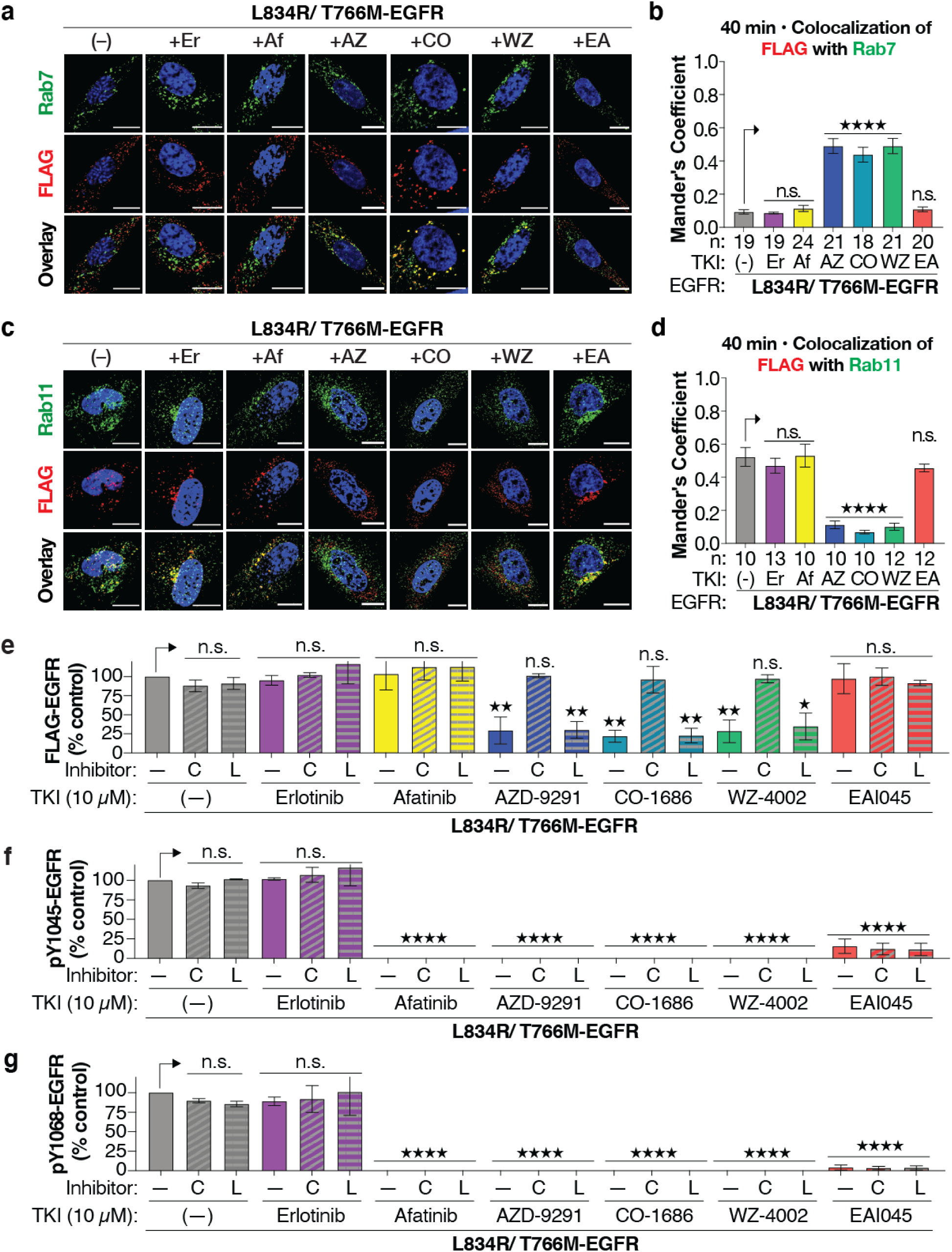
Clinically relevant, third-generation tyrosine kinase inhibitors influence L834R/T766M EGFR trafficking and induce EGFR degradation. Confocal microscopy of CHO-K1 cells expressing FLAG-tagged L834R/T766M EGFR (false colored red) and and immuno-labeled with **(a)** Rab7 (false colored green) as a marker for degradative endosomes or **(c)** Rab11 (false colored green) as a marker for recycling endosomes, 30 minutes after pre-incubation without/with 10 μM of indicated TKI. Scale bars = 10 µm. Bar plots illustrating the MCC value representing the colocalization of FLAG-tagged L834R/T766M EGFR with **(b)** Rab7 or **(d)** Rab11 without/with the indicated TKI. n = # of cells. Error bars = s.e.m. ****p<0.0001, ***p<0.001, **p<0.01, *p<0.1, n.s. not significant, from one-way ANOVA with Tukey’s multiple comparison test. **(e)** Normalized loss of FLAG-tagged L834R/T766M EGFR in CHO-K1 cells 12 hours following pre-incubation without/with 100 µM chloroquine (C) or lactacystin (L) (10 µM) and/or 10 µM Erlotinib (Er), afatinib (Af), AZD9291 (AZ), CO-1686 (CO), WZ-4002 (WZ), or EAI045 (EA). Phosphorylation of L834R/T766M EGFR at **(f)** Y1045; and **(g)** Y1068 in CHO-K1 cell lysates prepared as described in **5e**. In E,F,G, Error bars = s.e.m.. ****p<0.0001, ***p<0.001, **p<0.01, *p<0.1, n.s. not significant from one-way ANOVA with Dunnett’s multiple comparison test. See also Figure 5-figure supplement 1.

Finally we asked whether the change in endocytic trafficking induced by AZD-9291, CO-1686, and WZ-4002 also resulted in L834R/T766M EGFR degradation. We observed that treatment of L834R/T766M EGFR with AZD-9291, WZ-4002, and CO-1686 led to markedly (> 75%) reduced receptor levels after 12 hours when compared to untreated samples (Figure 5e, Figure 5–figure supplement 1b). In contrast, uninhibited, Afatinib-treated, and EAI045-treated L834R/T766M EGFR levels remained steady (Figure 5–figure supplement 1b). The TKI-induced degradation of L834R/T766M EGFR induced by AZD9291, CO-1686, and WZ-4002 was inhibited completely by chloroquine, while lactacystin had no effect (Figure 5e, Figure 5–figure supplement 1b). Neither chloroquine nor lactacystin affected the levels of L834R/T766M EGFR when mock-treated or in the presence of TKIs that failed to induce L834R/T766M EGFR degradation. For all cases inspected, predictable levels (Zhou et al., 2009; Walter et al., 2013; Cross et al., 2014; Jia et al., 2016) of phosphorylated L834R/T766M EGFR were detected at Y1045 (Figure 5f, Figure 5–figure supplement 1c) and Y1068 (Figure 5g, Figure 5–figure supplement 1d). Thus, third-generation TKIs that engage the ATP-binding pocket (Zhou et al., 2009; Walter et al., 2013; Cross et al., 2014), allosterically induce an EGF-type structure within the JM segment (Lowder et al., 2015), and traffic the receptor into Rab7+ endolysosomes (this work), also induce significant levels of L834R/T766M EGFR degradation through a lysosomal, as opposed to proteasomal mechanism.

## Discussion

EGFR is an essential, membrane-embedded sensor that communicates and integrates growth factor-dependent signals into diverse cellular phenotypes (E. Kovacs et al., 2015; Lemmon and Schlessinger, 2010). It is also one of the most potent oncogenes in human cancers (Pines et al., 2010; Sigismund et al., 2018) and remains an insufficiently addressed therapeutic target. EGFR is activated by growth factors that bind to the receptor extracellular domain (ECD) and by diverse mutations; in both cases the result is a dimeric receptor with one of two coiled coils within the cytoplasmic juxtamembrane segment (JM) and a catalytically active asymmetric kinase dimer. Despite its clear therapeutic significance, the mechanism by which EGFR decodes growth factor identity and/or mutational status into distinct and dynamic cellular signaling programs has remained elusive. In part, this knowledge gap results from the absence of a high-resolution view of how EGFR acts as an allosteric unit to communicate information across multiple domains and a complex lipid bilayer. But a more complete understanding of EGFR function is also precluded by allostery itself; how can one separate the activities of two or more protein domains whose conformational landscapes are dynamic and themselves tightly coupled?

### The JM coiled coil is a rotational toggle switch

In this work, we use structural, biochemical, and chemical biology tools to decouple the conformational landscape of the JM from both the extracellular domain and the kinase domain without loss of EGFR activity. Using these mutants and tools, we identify the cytoplasmic juxtamembrane segment as an essential EGFR processing center. The JM receives inputs from both the transmembrane helices and the kinase domain and assembles into one of two rotationally isomeric coiled coils. Together, these coiled coils contain all of the information necessary to specify both the direction of trafficking along the endocytic pathway and EGFR lifetime. These results support a model in which the pathway of endocytic trafficking following EGFR activation is determined by JM coiled coil status and not by growth factor-dependent differences in the EGFR-ligand complex stability as previously proposed (Ebner and Derynck, 1991; Francavilla et al., 2016; Roepstorff et al., 2009). The different effects of EGFR-selective growth factors also do not correlate with receptor dimer strength (Freed et al., 2017), as EGF and TGF-α both induce high affinity ECD dimers but traffic differently. The accompanying submission by Huang *et al*. complements these findings by providing a high-resolution view of how the binding of EGF and TGF-α to the extracellular domain alter the orientation of the dimeric receptor as it tracks into the membrane-embedded transmembrane helix. As changes in transmembrane helix orientation are known to bias JM conformation in cells (Sinclair et al., 2018), together the contributions provide a clear picture of how conformation changes within the EGFR extracellular domain are transduced into alternative JM conformations, and how alternative JM conformations are necessary and sufficient to control fundamental EGFR biology.

The structures of the EGF- and TGF-α-type coiled coils differ not only in the residues that mediate helix-helix interactions but also in the external surface available for intra- and intermolecular interaction. In the EGF-type structure, the residues at the *c*, *f*, and *b* positions of the coiled coil repeat are charged and polar(Jura et al., 2009), whereas these positions are hydrophobic (all leucine) in the TGF-α-type structure (Figure 1a). A sophisticated molecular dynamics-derived model of the active full-length receptor dimer (Arkhipov et al., 2013) shows the receiver kinase JM in the EGF-type structure with direct interactions between the receiver kinase JM latch (residues 664-681) and the charged surface of the activator kinase C-lobe. The details of these interactions and/or their dynamics would be different were the JM assembled into the TGF-α-type structure, whose interaction surface is hydrophobic, not polar and charged. The different interaction surfaces could interact differently with the kinase domain to alter the position (and thus accessibility) of the C-terminal tail, which is believed to localize near the JM (Jura et al., 2009; Brewer et al., 2009; Erika Kovacs et al., 2015) or alter the precise arrangement of acceptor and receiver kinase domains relative to one another (Arkhipov et al., 2013; Zhang et al., 2006; Wilson et al., 2009) and thereby influence their inherent substrate specificity. These different interaction surfaces could also play a direct role by differentially recruiting known JM-interacting factors such as, Nck adaptor protein (Hake et al., 2008), PKC (Hunter et al., 1984), p38MAPK (Winograd-Katz and Levitzki, 2006), AP2 (Kil et al., 1999; Huang et al., 2003) calmodulin (Viegas et al., 2020) and ARNO (Viegas et al., 2020), which in turn bias EGFR signaling and the consequent trafficking route. The presence of a LeuLeu motif on the outside surface of the TGF-α-type coiled coil is especially intriguing, as LeuLeu motifs elsewhere within EGFR (notably the JM-B region and C-terminal tail) are docking sites for endosomal sorting factors (Kil et al., 1999; Huang et al., 2003).

### Altered trafficking and degradation as kinase-independent EGFR and TKI activities

Small molecule EGFR inhibitors that bind near or within the ATP binding pocket and monoclonal antibodies targeting the extracellular domain have produced impressive therapeutic benefits to responsive cancers (Yuan et al., 2019). However, neither therapeutic modality offers sustained patient benefits, in large part because of acquired and innate resistance mechanisms (Thress et al., 2015; Westover et al., 2018). There is a growing realization that EGFR possesses kinase-independent pro-survival functions in cancer cells (Liu et al., 2018; Thomas and Weihua, 2019) that are insufficiently inhibited by either monoclonal antibodies or small molecules that inhibit kinase activity alone. Our results support a model in which pro-survival kinase-independent EGFR functions are related to JM-dependent differences in trafficking that avoid lysosomal degradation and receptor downregulation. Indeed, both EGFRvIII (implicated in glioblastoma multiforme) and L834R/T766M EGFR (implicated in NSCLC) avoid Cbl binding, ubiquitination, and degradation (Grandal et al., 2007; Shtiegman et al., 2007). In the case of L834R/T766M EGFR, this kinase-independent activity is reversed by TKIs that shift the JM coiled coil equilibrium to promote lysosomal trafficking and degradation. Identification of the as-yet-unknown factors that mediate this trafficking could serve as a new strategy for targeted protein degradation that complements both traditional PROTAC-like strategies and lysosomal targeting strategies based on cell surface receptor engagement (Ahn et al., 2021; Banik et al., 2020; Cotton et al., 2021).

## Materials and Methods

### Materials. Antibodies

Goat polyclonal anti-Rabbit, Horseradish Peroxidase (HRP)-conjugated (#7074), Goat polyclonal anti-Mouse, HRP-conjugated (#7076), Rabbit monoclonal anti-Phospho-EGF ReceptorTyr1173, (53A5) (#4407), Rabbit polyclonal anti-Phospho-EGF Receptor Tyr1086 (#2220), Rabbit polyclonal anti-Phospho-EGF Receptor Tyr1068 (#2234), Rabbit polyclonal anti-Phospho-EGF Receptor Tyr1045 (#2237), Rabbit monoclonal anti-α-Tubulin (#2125), Rabbit monoclonal anti-vinculin (#13901), Rabbit monoclonal anti-EEA1 (C45B10) (#3288), Rabbit monoclonal anti-Rab7 (D95F2) XP (#9367), Rab11 (D4F5) XP Rabbit mAb (#5589), Anti-mouse IgG (H+L) and F(ab’)2 Fragment Alexa Fluor® 555-conjugate (#4409) were purchased from Cell Signaling Technologies (CST). Mouse monoclonal (M2) anti-Flag (#F1804), Anti-FLAG® M2 Affinity Gel (#A2220) were purchased from Millipore Sigma. Goat anti-Mouse IgG (H+L) Cross-Adsorbed Secondary Antibody, Alexa Fluor 488®-conjugate (#A11001), IgG (H+L) Cross-Adsorbed Goat anti-Rabbit, Alexa Fluor™ 488 (#A11008), were purchased from ThermoFisher Scientific.

### Chemicals and Recombinant Proteins

F-12K Medium (#10-025-CV), Dulbecco’s Phosphate Buffered Saline (DPBS) (#14190), Fetal Bovine Serum (FBS)–Heat Inactivated (#11082147), Penicillin/Streptomycin (#1514012), Non-enzymatic Cell Dissociation Solution (#13151014), RestoreTM Western Blot Stripping Buffer (#21059), Hoechst 33342, Trihydrochloride, Trihydrate - 10 mg/mL Solution in Water (#H3570), iBlot PVDF membranes (# IB401031) were purchased from ThermoFisher Scientific. FugeneHD transfection reagent (E2311) was purchased from (Promega). Fetal Bovine Serum (FBS)–Heat Inactivated (#F4135), Bovine Serum Albumin (#9048-46-8), Fibronectin (#F1141) were purchased from Millipore Sigma. cOmplete, Mini Protease Inhibitor Tablets (#11836170001), PhosSTOP Phosphatase Inhibitor Cocktail Tablets (#04906837001) were purchased from Roche Applied Science. Recombinant Human EGF Protein (#236-EG), Recombinant Human TGF-a Protein (#293-A) were purchased from R&D Systems. Mini-PROTEAN® TGXTM Precast Gels (10% polyacrylamide) (#456-1036), ClarityTM Western ECL reagents (#1705060) were purchased from Bio-Rad Laboratories, Inc. Lactacystin, proteasome inhibitor (#ab141411) and Chloroquine diphosphate, apoptosis and autophagy inhibitor (#ab142116) were purchased from AbCam. Tyrosine Kinase Inhibitors (TKIs) Erlotinib HCl (OSI-744) (#S1023), Afatinib (BIBW2992) (#S1011), Osimertinib (AZD-9291) (#S7297), Rociletinib (CO-1686) (#S7284), WZ4002 (#S1173), EAI045 (#S8242) were purchased from Selleck Chemicals.

### Cell culture

CHO-K1 cells (ATCC) were cultured in F12K Medium supplemented with 10% FBS and Pen-Strep (100 I.U./mL penicillin and 100 mg/mL streptomycin) at 37°C in a CO2/air (5%/95%) incubator. Cells were transfected using the TransIT-CHO Transfection Kit (Mirus Bio LLC) (CHO-K1) or using FugeneHD (Promega), according to the manufacturer’s instructions. Cell densities for all mammalian cell lines were determined with a Cellometer® Auto T4 automated counter. All cells were bona fide lines and periodically tested for mycoplasma with DNA methods

### Cloning and Mutagenesis

All EGFR DNA variants were cloned from a pcDNA3.1 plasmid, generously donated by the Kuriyan Group (University of California, Berkeley), containing the sequence of the full-length EGFR with an N-terminal FLAG tag14. Mutations were introduced into the wild-type, CCH-1 and CCH-10 EGFR sequences using Quikchange Lightning site-directed mutagenesis kit (Agilent Technologies), according to the manufacturer’s instructions, with primers (purchased from Integrated DNA Technologies) listed in Figure 1– Supplemental Table 1. All DNA variants were amplified with XL-10 Gold Ultracompetent cells (Agilent Technologies).

### Bipartite Tetracysteine Display Assay i.e. Surface ReAsH Labeling Studies and Total Internal Resonance Fluorescence (TIRF) Microscopy

ReAsH labeling was accomplished as described previously(Doerner et al., 2015; Sinclair et al., 2018) by treating CHO-K1 cells expressing EGFR variants with an endocytosis inhibition cocktail (10 mM NaN3, 2 mM NaF, 5 mM 2-deoxy-D-glucose in F12-K media) for 1 hr at 37°C. Cells were stimulated without/with 100 ng/mL of EGF (16.7 nM)) and TGF-α (16.7 nM) prior to labeling. Cells were washed once with endocytosis inhibitor-containing media before incubation with ReAsH labeling solution (2 mM ReAsH (ThermoFisher Scientific), 20 mM BAL (Acros Organics) in F12K media) for 1 hr at 37°C. Cells were washed and incubated with endocytosis inhibitor-containing F12K media supplemented with 100 mM BAL for 10 min at 37°C. The media was removed, and cells fixed using 4% paraformaldehyde (PFA) in DPBS for 25 min at room temperature. Fixed cells were washed with DPBS and blocked with 10% BSA in DPBS for 30 min at 37°C. Cells were then labeled with primary antibody (mouse monoclonal mouse M2 anti-FLAG, 1:1000 dilution in 10% BSA in DPBS) for 1 hr at 37°C, washed thrice with 10% BSA in DPBS, then incubated with secondary antibody (AlexaFluor488-conjugated goat anti-mouse, 1:2000 dilution in 10% BSA in DPBS) for 1 hr at 37°C. Cells were then washed twice with 10% BSA in DPBS, washed once with DPBS, then nuclear-stained with Hoescht 33342 (1.62 mM in DPBS) for 5 min at 37°C. Cells were then washed once with DPBS and stored in DPBS at 4°C, prior to imaging. Labeled cells were monitored via TIRF microscopy, conducted on a Leica microsystems AM TIRF MC DMI6000B fitted with an EM-CCD camera (Hamamatsu) with HCX PL APO 63x/1.47 oil corrective objectives, as described previously(Doerner et al., 2015; Sinclair et al., 2018). Images were analyzed with ImageJ (FIJI) as described previously(Doerner et al., 2015; Sinclair et al., 2018).

### Immunofluorescent labeling, Confocal microscopy and Image analysis

Immunofluorescent labeling and confocal microscopy to assess localization of EGFR variants expressed in CHO-K1 cells was carried out as described previously(Francavilla et al., 2016) with slight modification. CHO-K1 cells expressing FLAG-tagged EGFR variants were incubated without/with 100 ng/mL of EGF (16.7 nM) or TGF-α (16.7 nM) for 1 hr at 4°C to allow growth factor binding. For experiments with TKIs, instead of growth factor treatment, cells were incubated with 10uM TKI (as indicated) in F12K medium at 4°C to allow TKI binding. Cells were then washed with DPBS and incubated with serum free media at 37°C for 8 or 40 minutes as indicated to allow endocytosis. Cells were then fixed using 4% PFA in DPBS for 25 min at room temperature. Cells were washed with DPBS and incubated with blocking buffer (5% normal goat serum (CST), 0.3% Triton X-100 in DPBS) for 1 hr at 37°C. Cells were then labeled with indicated primary antibodies overnight (∼12 hrs) at 4°C (mouse M2 anti-Flag, 1:1000 dilution and rabbit anti-Rab7, 1:1000 dilution or rabbit anti-EEA1, 1:1000 dilution or rabbit anti-Rab11, 1:1000 dilution in antibody dilution buffer (1% BSA, 0.3 % Triton X-100 in DPBS)). Cells were then washed thrice with DPBS and incubated with secondary antibody (AlexaFluor488-conjugated goat anti-rabbit, 1:500 dilution or AlexaFluor555-conjugated goat anti-mouse, 1:500 dilution in antibody dilution buffer) for 2 hr at room temperature. Cells were then washed twice with DPBS and nuclear-stained with Hoescht 33342 (1.62 mM in DPBS) for 5 min at room temperature. Cells were then washed once with DPBS and stored in DPBS at 4°C, prior to imaging. Laser-scanning confocal microscopy experiments of labeled immunofluorescent samples were performed at room temperature on an inverted Zeiss LSM 880 laser-scanning confocal microscope equipped with a Plan-Apochromat ×63/1.4 numerical aperture oil immersion lens and a diode 405 nm laser, an Argon 458, 488, 514 nm laser, a diode pumped solid-state 561 nm laser and a 633 nm HeNe laser with standard settings. DAPI and Alexa-488, Alexa Fluor 555, and Alexa Fluor 647 dyes were excited with 405-, 488-, 546-, and 633-nm laser lines, and emitted light was collected through band pass filters transmitting wavelengths of 420–480 nm, 505–530 nm and 560–615 nm and a long-pass filter transmitting 615 nm, respectively. The pinhole size was set to 1 airy unit. Images were acquired at a nuclear section with fixed thresholds. Image acquisition was performed with ZEN software (Carl Zeiss). Raw images were exported as .lsm files. Images were analyzed using ImageJ software(Schneider et al., 2012). Colocalization of EGFR with indicated endocytic marker (Rab7, Rab11, EEA1) was evaluated as Manders’ Colocalization Coefficient (MCC)(Bolte and Cordelières, 2006) which represents the sum of intensities of green pixels (due to Rab11 or Rab7 or EEA1) that also contain red (due to FLAG-tagged EGFR) divided by the total sum of green intensities. Colocalization was evaluated using JACoP (Just Another Colocalization Plugin)(Dunn et al., 2011) in ImageJ. MCC values for each condition obtained from multiple cells collected over at least 2 biological replicates were pooled and represented as Mean with S.E.M using Prism 8.4.3

### Western Blot Analysis of EGFR Expression and Autophosphorylation

Western blot analysis of EGFR expression and autophosphorylation in transfected CHO-K1 cells was accomplished as described previously with slight modification (Doerner et al., 2015; Sinclair et al., 2018). CHO-K1 cells expressing FLAG-tagged EGFR variants were serum starved overnight (∼12 hours). 48 hr post seeding cells were stimulated without/with 100 ng/mL of EGF (16.7 nM) or TGF-α (16.7 nM) for 5 min at 37°C, washed with serum free F12K media, and lysed in 100 mL of lysis buffer (50 mM Tris,150 mM NaCl, 1 mM EDTA, 1 mM NaF, 1% Triton X-100, pH 7.5, 1x cOmplete protease inhibitor cocktail, 1x Phos-Stop) for 1 hr. For experiments investigating the time dependent changes in intracellular EGFR levels, CHO-K1 cells were incubated without/with 100 ng/mL of EGF (16.7 nM) or TGF-α (16.7 nM) for 15 min following which, growth factor solution was washed with DPBS and cells were incubated at 37°C with serum free media for 0-90 minutes and cell lysis was carried out as described previously. For experiments investigating the time dependent changes in EGFR phosphorylation, CHO-K1 cells were incubated without/with 100 ng/mL of EGF (16.7 nM) or TGF-α (16.7 nM) for 15 min following which, growth factor solution was washed with DPBS and cells were incubated at 37°C with serum free media for 0-8 minutes and cell lysis was carried out as described previously. Clarified cell lysates were subjected to reducing 4-15% polyacrylamide SDS-PAGE electrophoresis and transferred to immuno-blot PVDF membranes. Membranes were blocked with 5% milk in TBS-T Buffer (50 mM Tris, 150 mM NaCl, 0.1% Tween, pH 7.4) for 1 hr followed by an overnight incubation at 4°C of indicated primary (rabbit or mouse) antibodies. Blots were washed with TBS-T and incubated with either anti-rabbit or anti-mouse goat horseradish peroxidase conjugate secondary antibodies for 1 hr at room temperature, then washed with TBS-T. Blots were then visualized using Clarity Western ECL reagents on a ChemiDoc XRS+/ ChemiDocMP instrument, and intensities of immuno-stained bands measured with ImageJ 64(Schneider et al., 2012). When assessing phosphorylation of EGFR/ gel loading at multiple positions using the same samples, the blots obtained with a given phospho-EGFR antibody were stripped with Restore Western Blot Stripping Buffer/ and antibody stripping buffer (Tris-HCl (62.5 mM), SDS (2%w/v), 2-mercaptoethanol (0.7%v/v)) and re-probed with a different phospho-EGFR antibody. For experiments investigating the time dependent changes in intracellular EGFR levels, the FLAG signal (total EGFR) was normalized to the vinculin loading control and normalized signal for the condition without any growth factor/ inhibitor treatment. For experiments investigating the time dependent changes in EGFR phosphorylation, the phospho-EGFR signal (pY1045, pY1068, pY1173) was normalized to the total EGFR (FLAG signal) and vinculin/ tubulin was used as a loading control. For western blot experiments with the tyrosine kinase inhibitors (TKI), CHO-K1 cells expressing L834R/ T766M EGFR were pre-treated with serum free F12K medium containing 10uM of TKI at 37°C for 30 minutes, washed once with DPBS followed by incubation with serum free DMEM for 12 hrs, prior to cell lysis. FLAG, pY1045 and pY1068 signal was normalized to the vinculin loading control and corresponding signal detected at 12 hours without inhibitor/ TKI treatment. For all experiments with proteasomal/ lysosomal inhibitors, the experiments were carried out with CHO-K1 cells as described before in the methods, involving a pretreatment with 10uM Lactacystin or 100uM Chloroquine for 1 hr at 37°C prior to growth factor or TKI treatment.

## Data availability

The materials and data reported in this study are available upon reasonable request from the corresponding author.

## Acknowledgements

This work was supported by the NIH (RO1 GM83257 and R35 GM134963 to A.S.). D.M. acknowledges training support provided by NIH (5T32GM008283-30). A.D. acknowledges training support provided by NIH (5T32GM008283-28). S.H.H.C. acknowledge the support provided by the Ministry of Education of Taiwan.

## Author Contributions

D.M., S.H.H.C, K.Q., A.D., and A.S. designed experiments. D.M., S.H.H.C, K.Q., and A.D. performed experiments. D.M., S.H.H.C, K.Q., A.D. and A.S. wrote the paper.

## Additional information

The figure supplements for the main text of this paper are available in an accompanying document.

## Supplementary Information

### List of Supplemental Figures

**Figure 1–figure supplement 1:**
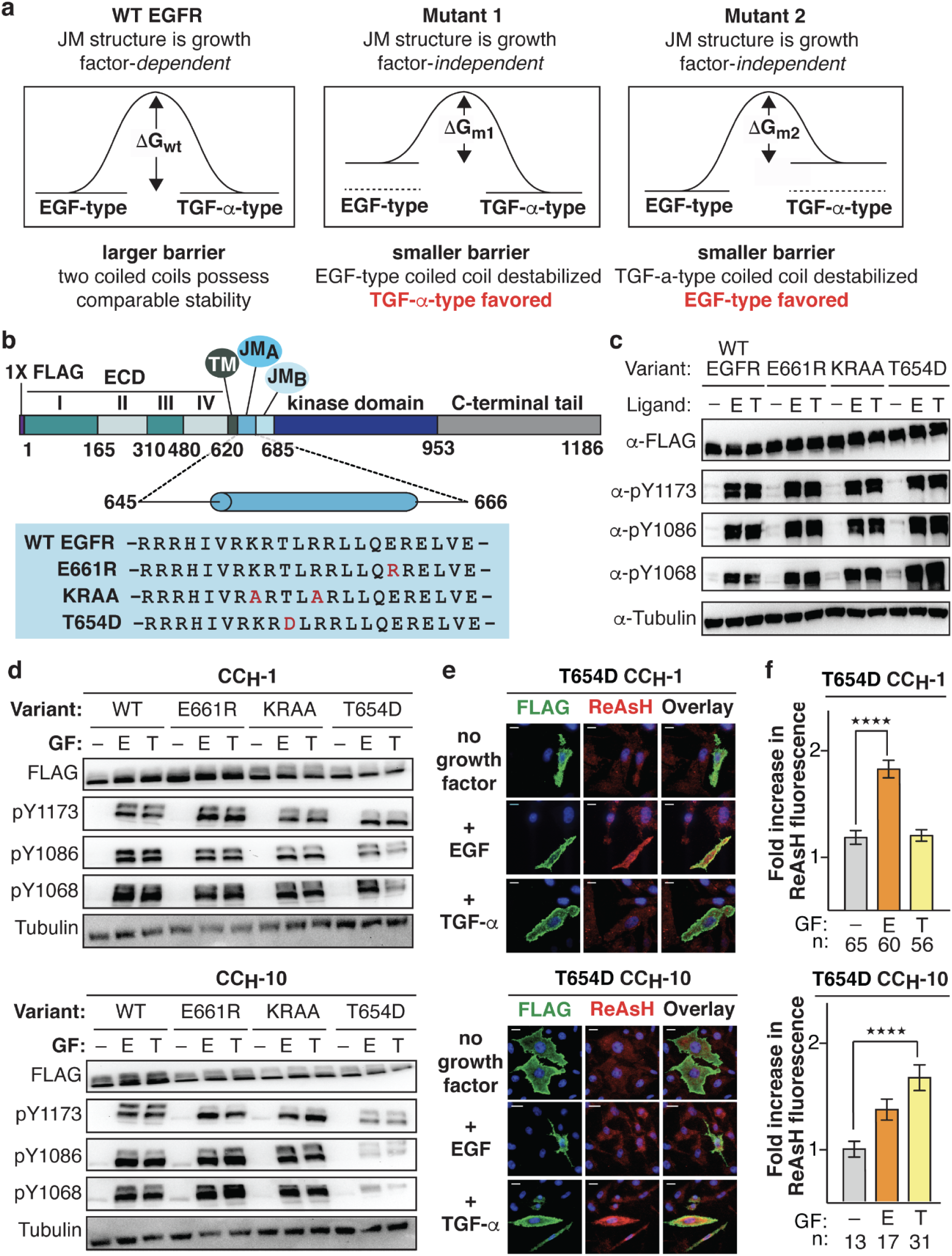
Design of EGFR decoupling mutants (KRAA, E661R) and controls (T654D) and TIRF microscopy images and western blots related to Bipartite tetracysteine display experiments. **(a)** Influencing coiled coil preferences by design. In WT EGFR, both coiled coil conformations are energetically accessible, and the identity of the bound growth factor influences which conformation is adopted. In Mutant 1, the EGF-type coiled coil is destabilized and the TGF-α-type structure is favored; in Mutant 2, the TGF-α-type coiled coil is destabilized and the EGF-type structure is favored. **(b)** Domain diagram of FLAG-tagged EGFR illustrating sequences of WT EGFR as well as E661R and KRAA decoupling mutants that favor the EGF-type or TGF-α-type coiled coil, respectively; T654D EGFR also contains a mutation within the JM but at a location predicted to not affect relative coiled coil stability (light blue box). **c,** E661R, KRAA, and T654D EGFR respond like WT EGFR to growth factor stimulation. Representative western blots illustrating expression and extent of Y1173, Y1086, and Y1068 phosphorylation of **(c)** FLAG-tagged WT, E661R, KRAA, and T654D EGFR and **(d)** FLAG-tagged CCH-1 and CCH-10 variants of WT, E661R, KRAA, and T654D EGFR in CHO-K1 cells stimulated continuously without/with EGF and TGF-α (16.7 nM) for 5 minutes 37°C. Alpha-tubulin is used as loading control. **(e)** Representative TIRF-M images of CHO-K1 cells illustrating ReAsH labeling (red fluorescence) and expression (green fluorescence) of FLAG-tagged CCH-1 and CCH-10 variants of T654D without/ with EGF/ TGF-α stimulation (16.7 nM). Scale bars = 10 µm. T654D EGFR was prepared as a control: this mutation occupies the *c* or *e* position of a coiled coil repeat when the JM assembles into the EGF- and TGF-α-type structures, respectively. **(f)** Bar plots illustrating the quantification of TIRF-M results from ‘n’ cells expressing FLAG-tagged T654D EGFR as fold increase in expression-corrected ReAsH fluorescence over background. Error bars = s.e.m. ****p<0.0001, n.s. p>0.05 from one-way ANOVA with Dunnett’s post-analysis accounting for comparisons within individual mutants to no growth factor treated control. n.s., not significant. Scale bars = 10 µm.

**Figure 2–figure supplement 1:**
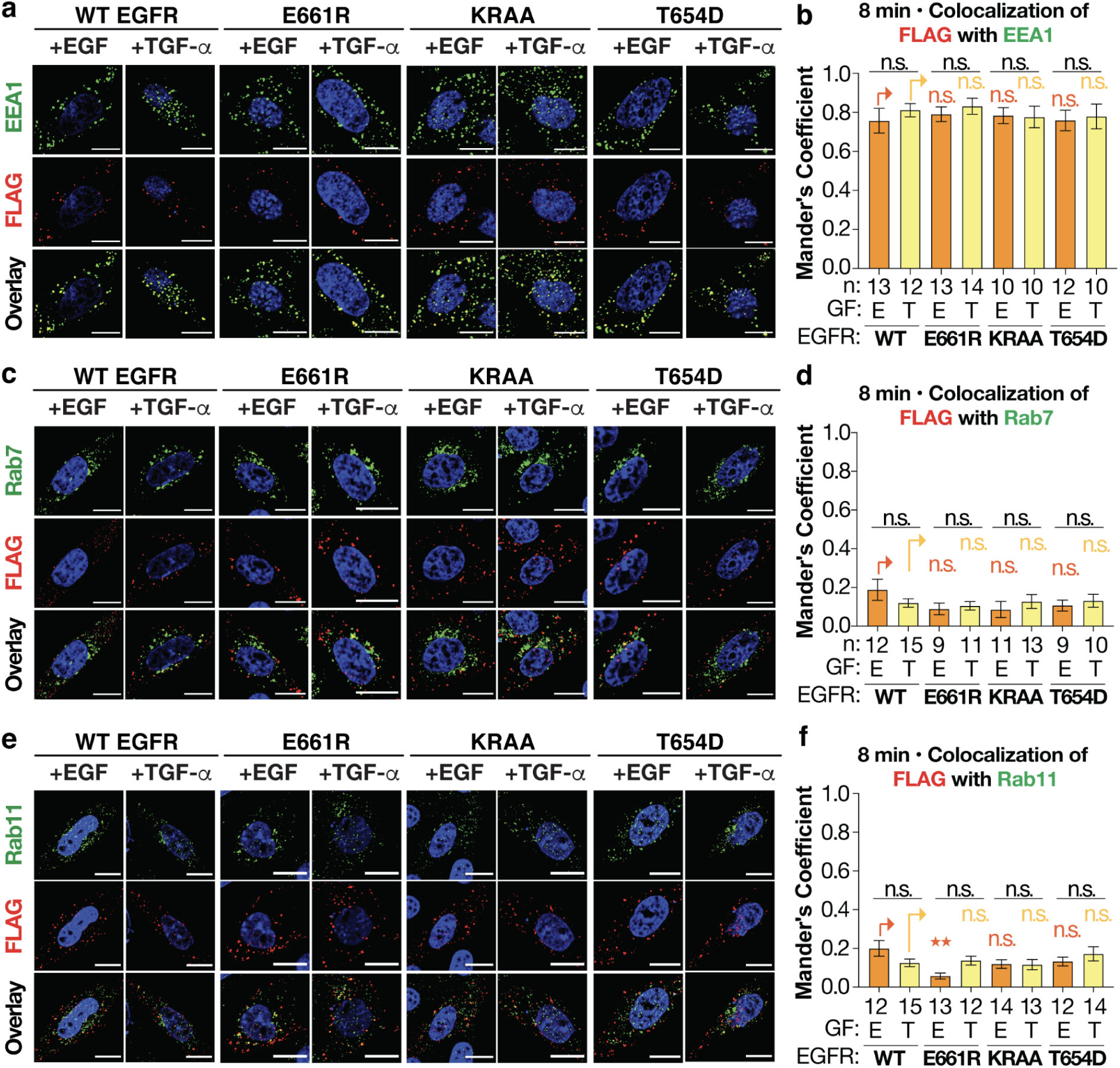
FLAG-tagged WT, E661R, and KRAA EGFR colocalize with EEA1 and not with Rab7 or Rab11 respectively, 8 minutes after stimulation with EGF or TGF-α. **(a, c, e)** Representative confocal microscopy images of CHO-K1 cells expressing FLAG-tagged WT, E661R, KRAA and T654D EGFR (false-colored red) 8 minutes after stimulation with EGF or TGF-α (16.7 nM). **(a)** Early endosomes (false-colored green) are identified using anti-EEA1 antibody. **(c)** Degradative endosomes (false-colored green) are identified using an anti-Rab7 antibody. **(e)** Recycling endosomes (false-colored green) are identified using an anti-Rab11 antibody. (**a, c, e)** Scale bars = 10 µm. Bar plots illustrating the quantified Mander’s co-localization coefficient (MCC) values of FLAG-tagged WT, E661R, KRAA and T654D EGFR with **(b)** EEA1 **(d)** Rab7 **(f)** Rab11 8 minutes after stimulation with EGF/ TGF-α (16.7 nM) for ‘n’ cells. Error bars, s.e.m., n.s. not-significant from one-way ANOVA with Tukey’s multiple comparison test.

**Figure 2–figure supplement 2:**
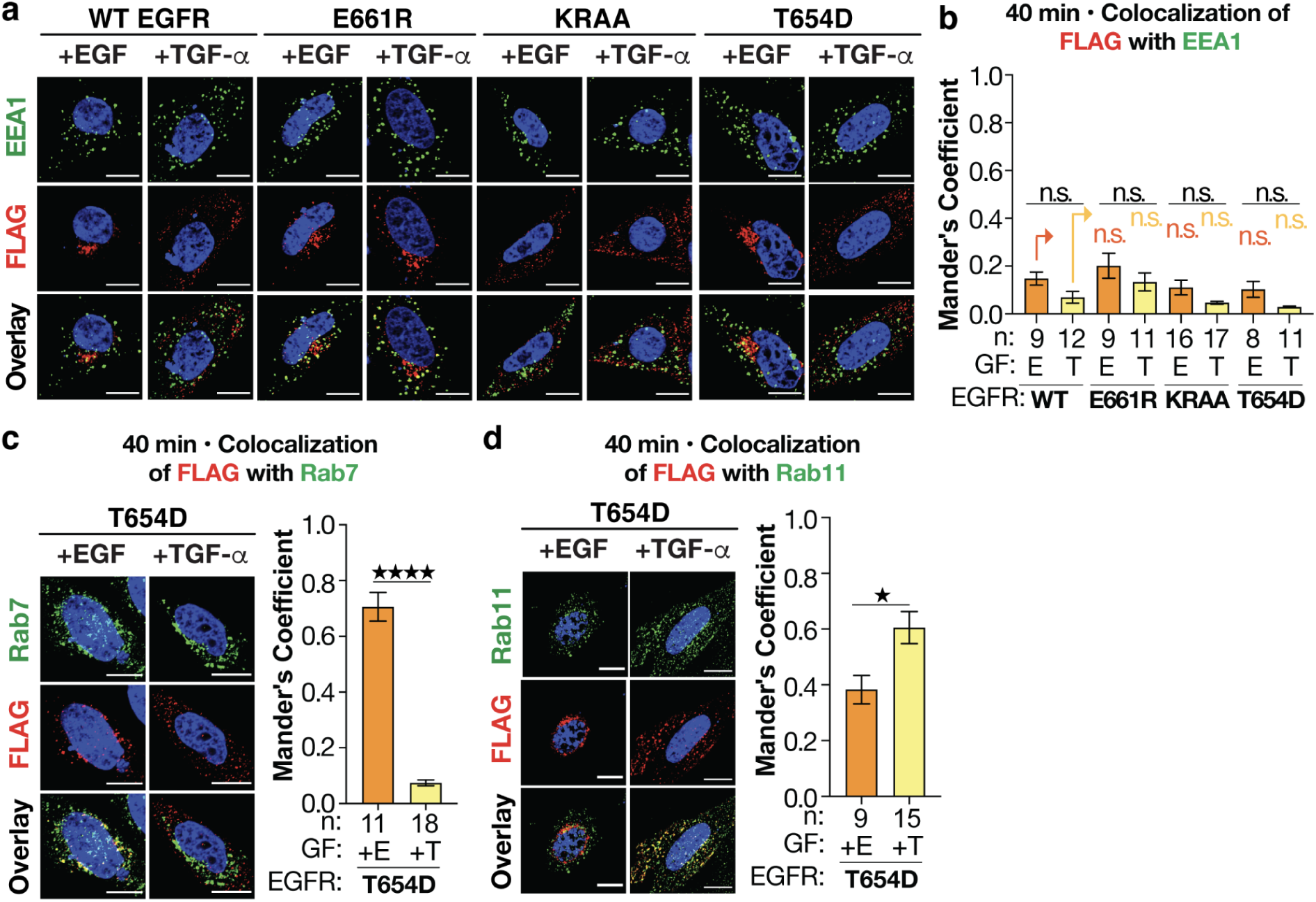
Co-localization of FLAG-tagged WT, E661R, KRAA and T654D EGFR with EEA1, 40 minutes after stimulation with EGF/TGF-α and co-localization of FLAG-tagged T654D EGFR with Rab7, Rab11, 40 minutes after stimulation with EGF/TGF-α. **(a)** Representative confocal microscopy images of CHO-K1 cells expressing FLAG-tagged WT, E661R, KRAA and T654D EGFR (false colored red) 40 minutes after stimulation with EGF or TGF-α (16.7 nM). Early endosomes (false colored green) are identified using an anti-EEA1 antibody. Scale bars = 10 µm. **(b)** Bar plots illustrating the quantified MCC values of FLAG-tagged WT, E661R, KRAA and T654D EGFR with EEA1 40 minutes after stimulation with EGF or TGF-α (16.7 nM) for ‘n’ cells. Error bars, s.e.m., n.s. not significant from one-way ANOVA with Tukey’s multiple comparison test. **(c, d)** Confocal microscopy of CHO-K1 cells expressing FLAG-tagged T654D EGFR (false-colored red) 40 minutes after stimulation with EGF/ TGF-α. **(c)** Degradative endosomes are identified using an anti-Rab7 antibody (false-colored green). **(d)** Recycling endosomes are identified using an anti-Rab11 antibody (false-colored green). Scale bars = 10 µm. Bar plots illustrating the quantified MCC values of FLAG-tagged T654D (green) with either **(c)** Rab7 or **(d)** Rab11 40 minutes after stimulation with EGF or TGF-α (16.7 nM) for ‘n’ cells. Error bars, s.e.m. ****p<0.0001, *p<0.1, from t-test.

**Figure 2–figure supplement 3:**
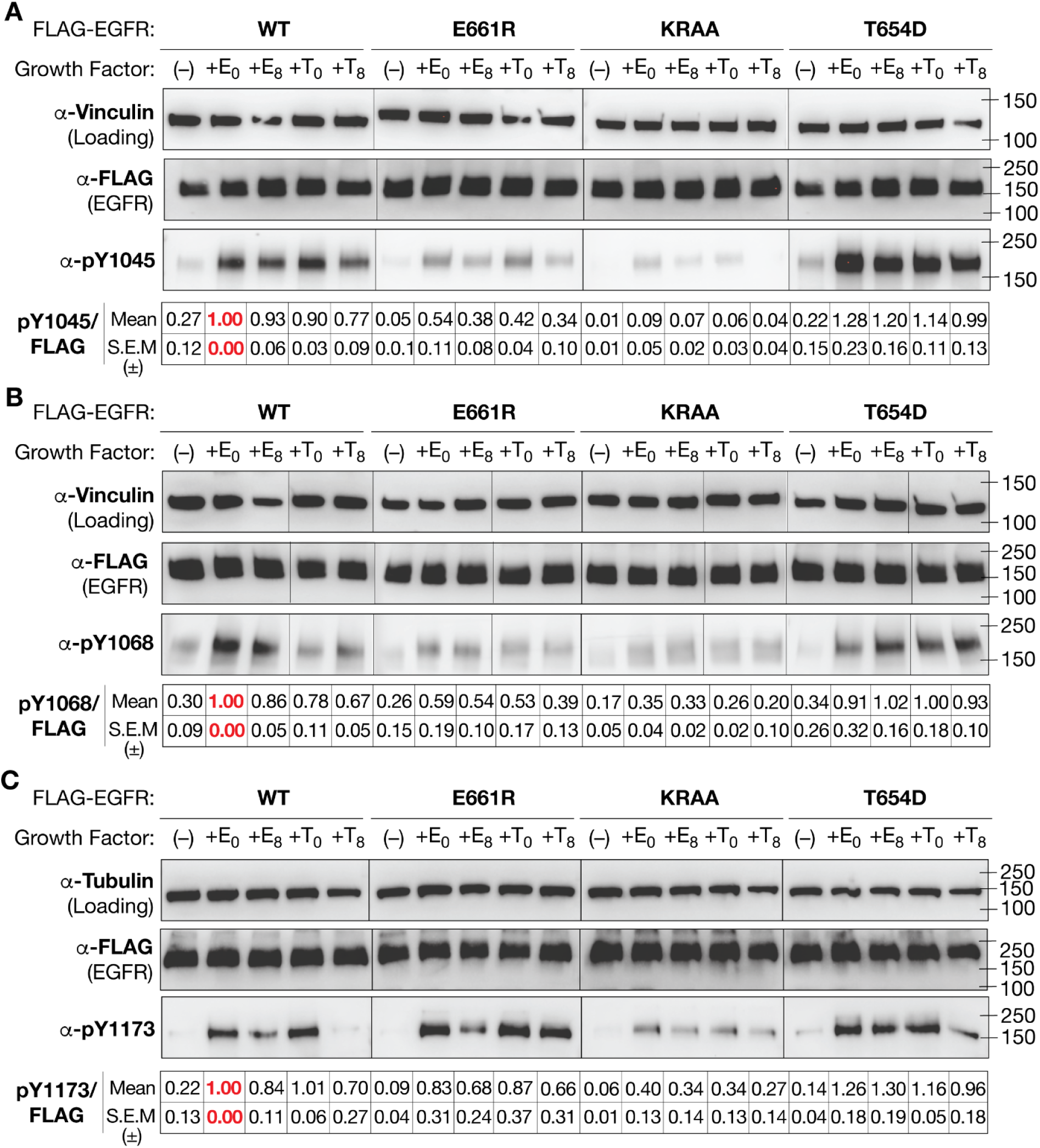
Time dependent decay in phosphorylation of FLAG-tagged WT, E661R, KRAA and T654D EGFR at Y1045, Y1068, Y1173. Immunoblots illustrating the time-dependent change in (A) pY1045; (B) pY1068; and (C) pY1173 0-8 minutes after stimulation with EGF or TGF-α (16.7 nM). FLAG blot indicates levels of total EGFR. Vinculin is used as a loading control. Blots represent data pooled from at least 3 biological replicates and densitometric quantification of signal pYEGFR/ FLAG (Mean and S.E.M.) is included below the immunoblots.

**Figure 3–figure supplement 1:**
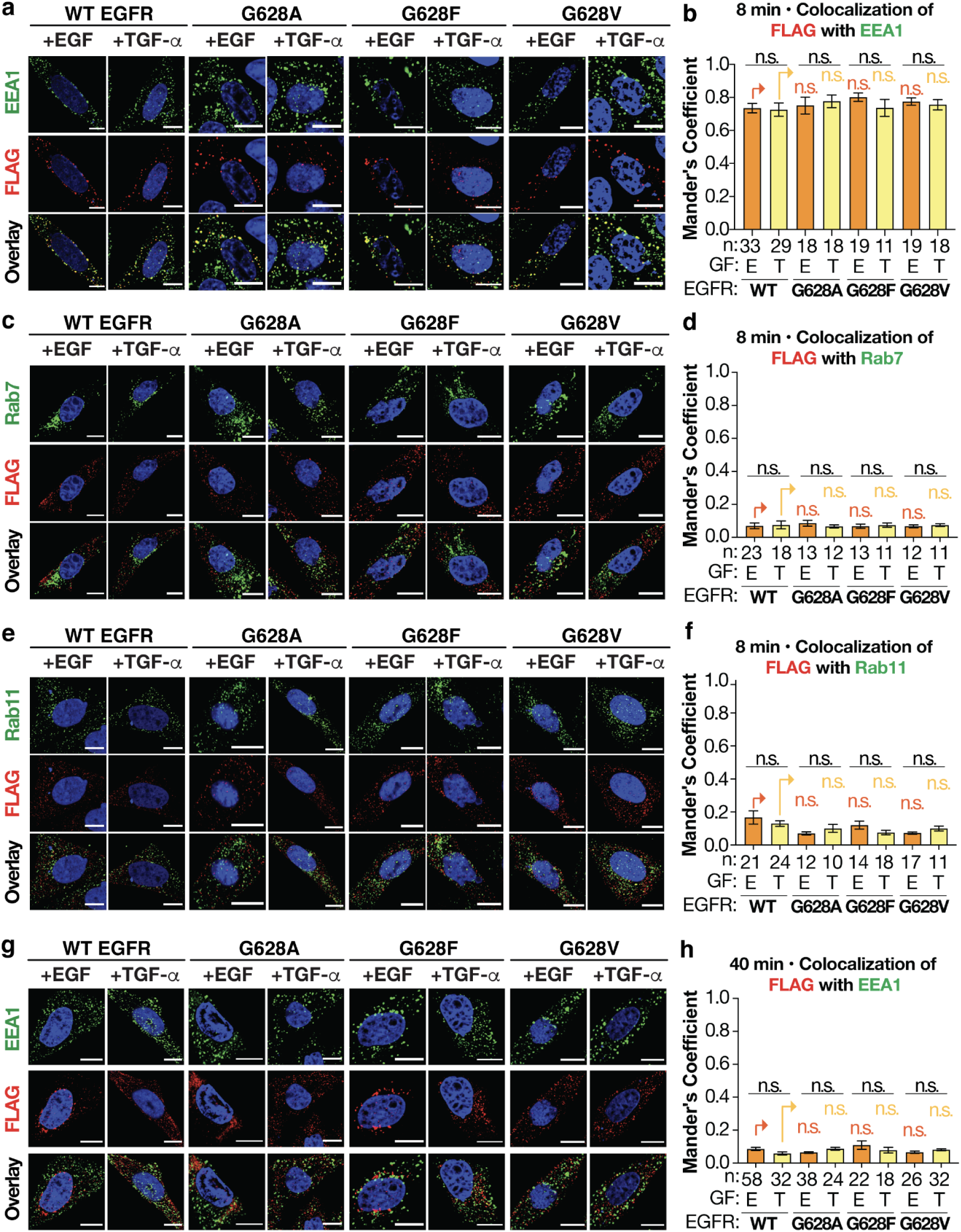
Co-localization of FLAG-tagged G628A, G628F, and G628V EGFR with EEA1 (8 and 40 min) and Rab7 or Rab11 (8 min) after stimulation with EGF or TGF-α. Representative confocal microscopy images of CHO-K1 cells expressing FLAG-tagged WT, G628A, G628F, and G628V EGFR (false colored red) stimulated for (**a, c, e)** 8 minutes; and **(g)** 40 minutes with EGF/ TGF-α (16.7 nM). (**a, g)** Early endosomes (false-colored green) are identified using an anti-EEA1 antibody. (**c)** Degradative endosomes (false-colored green) are identified using an anti-Rab7 antibody. (**e)** Recycling endosomes (false-colored green) are identified using an anti-Rab11 antibody. (**a, c, e, g)** Scale bars = 10 µm. Bar plots illustrating the quantified MCC values of FLAG-tagged WT, G628A, G628F, G628V EGFR with (**b)** EEA1, 8 min or **(d)** Rab7 or **(f)** Rab11 or **(h)** EEA1, 40 min; after stimulation with EGF/ TGF-α (16.7 nM) for ‘n’ cells. Error bars, s.e.m., n.s. not significant from one-way ANOVA with Tukey’s multiple comparison test.

**Figure 3–Figure supplement 2:**
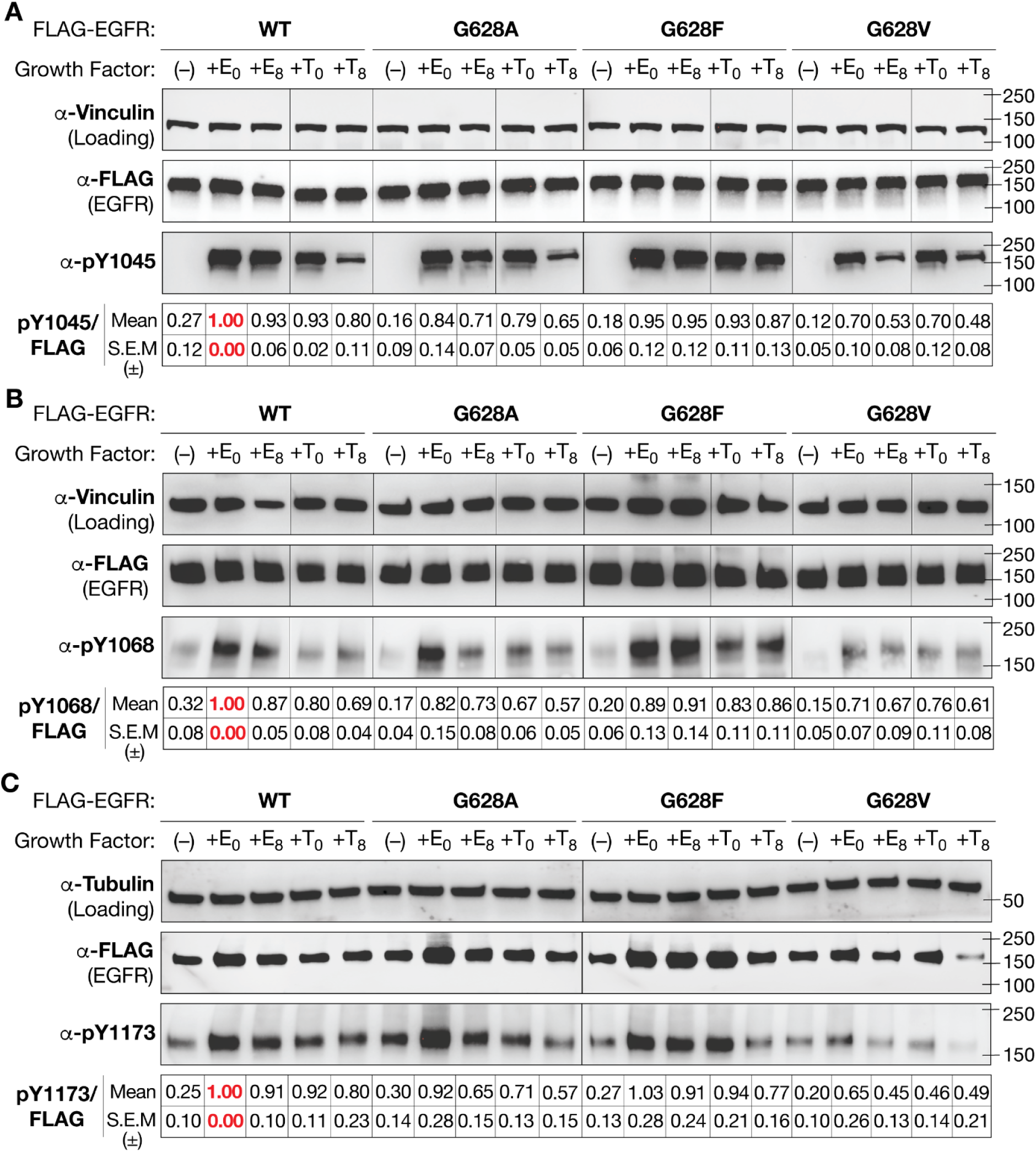
Time dependent decay in phosphorylation of FLAG-tagged WT, G628A, G628F and G628V EGFR at Y1045, Y1068, Y1173. Immunoblots illustrating the time-dependent change in **(a)** pY1045; **(b)** pY1068; and **(c)** pY1173 for FLAG-tagged WT EGFR, G628A, G628F and G628V 0-8 minutes after stimulation with EGF or TGF-α (16.7 nM). FLAG blot indicates levels of total EGFR. Vinculin/ Alpha-Tubulin is used as a loading control. Blots represent data pooled from at least 3 biological replicates and densitometric quantification of pYEGFR/FLAG signal (Mean and S.E.M.) is included below the immunoblots.

**Figure 5–figure supplement 1.**
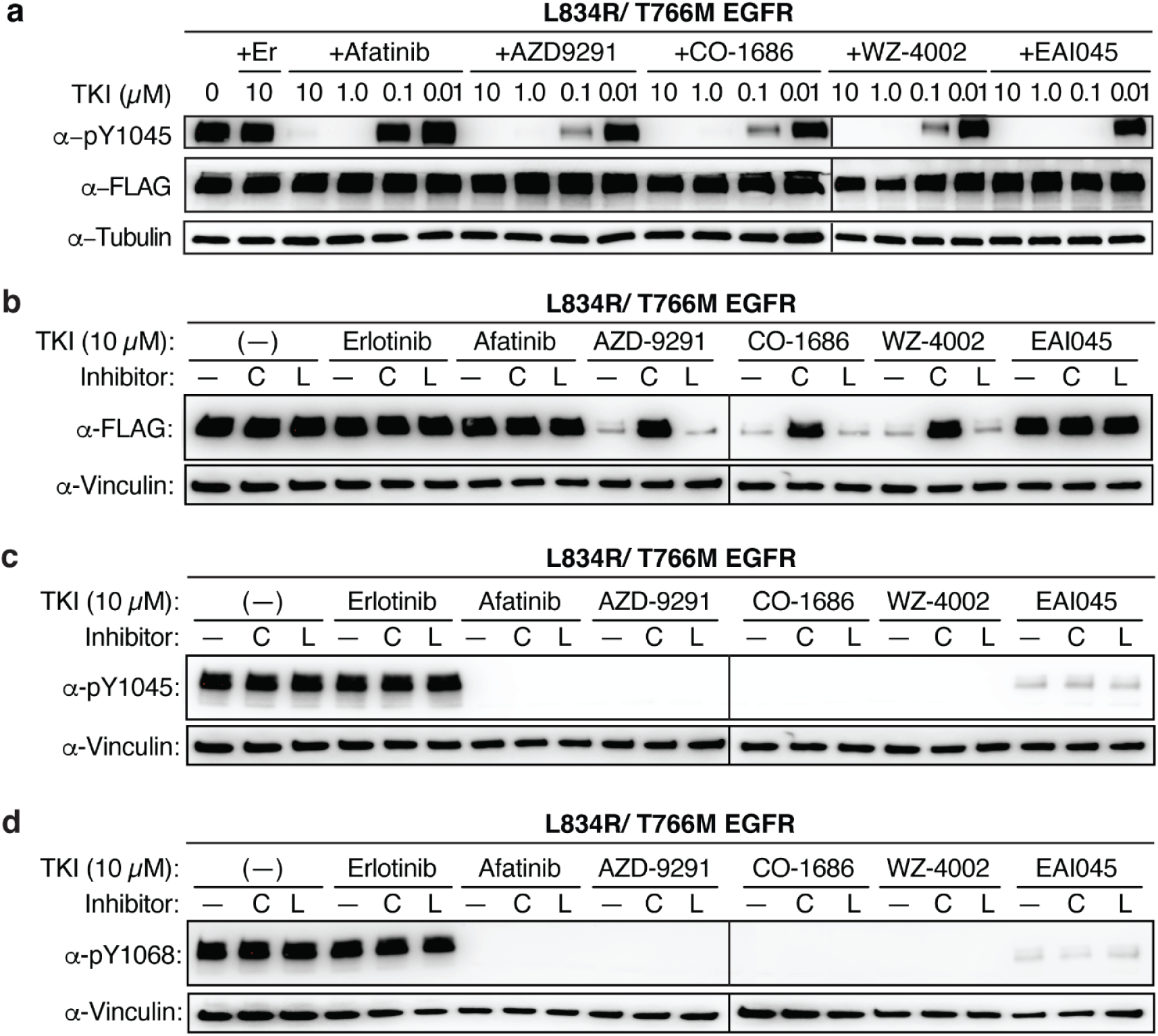
Immunoblots illustrating the intracellular levels of FLAG-tagged and phosphorylated EGFR detected at various time points post treatment with EGFR TKIs. **(a)** Immunoblots illustrating the cellular expression (FLAG) and dose-dependent phosphorylation of FLAG-tagged L858R/T790M EGFR at Y1045, detected in CHO-K1 cells 30 minutes following pre-incubation without/ with 10, 1.0, 0.1 and 0.01 uM of TKIs erlotinib (+Er), afatinib, AZD9291, CO-1686, WZ-4002 and EAI045. Alpha-tubulin is used as loading control. **(b)** Immunoblots illustrating the loss of FLAG-tagged L834R/T766M EGFR detected in CHO-K1 cells lysed 12 hours following pre-incubation without/with 100 µM chloroquine, C or lactacystin, L (10 µM) for 1 hour followed by pre-incubation without/with 10 µM erlotinib, afatinib, AZD9291, CO-1686, WZ-4002, or EAI045. Immunoblots illustrating the phosphorylation of L834R/T766M EGFR at **(c)** Y1045; and **(d)** Y1068 in CHO-K1 cell lysates prepared as described in **c**. Vinculin is used as loading control.

**Supplemental Table 1.**
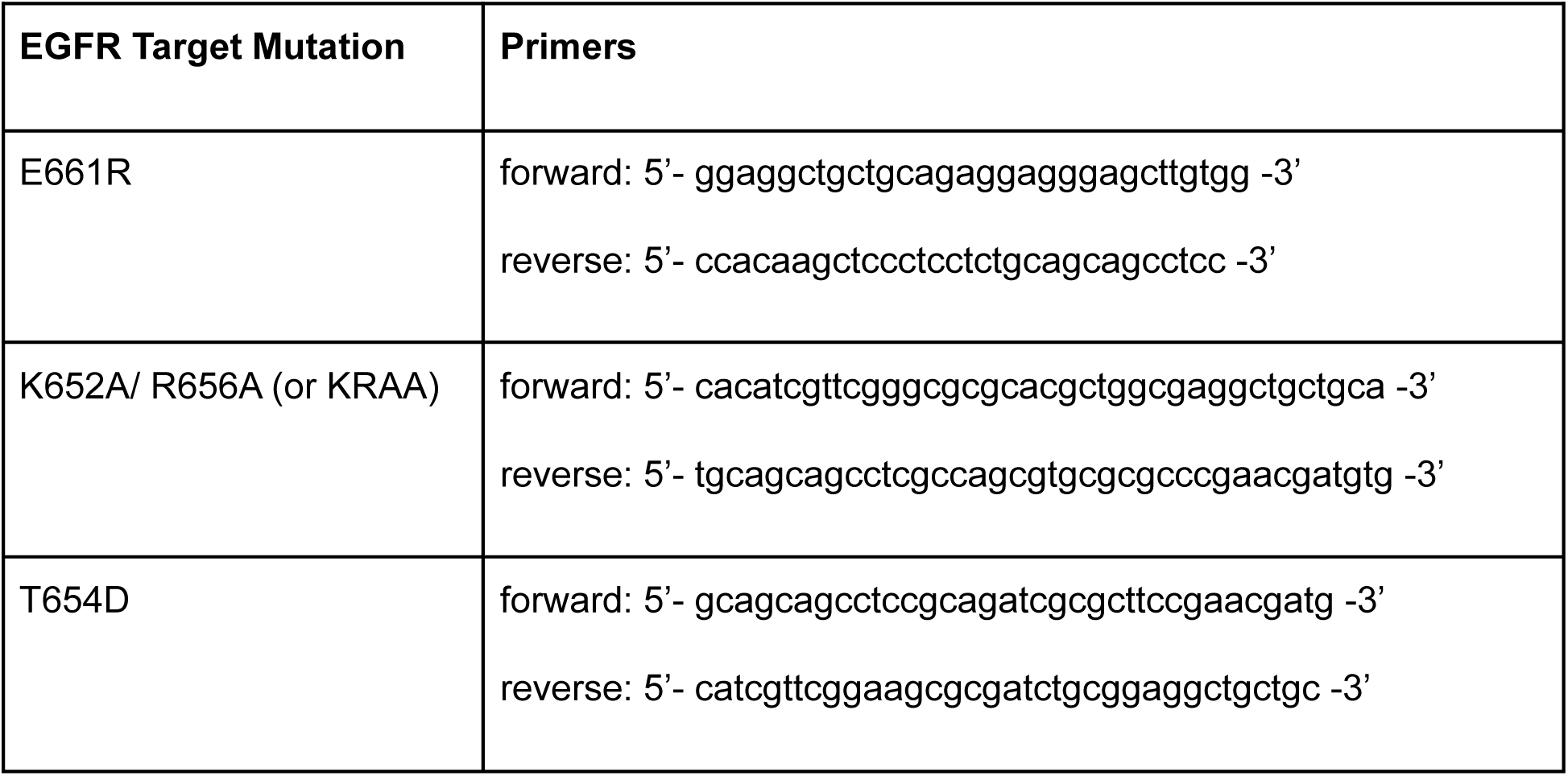
List of mutagenesis primers used to design JM decoupling mutants

